# Evaluation of tick salivary and midgut extracellular vesicles as anti-tick vaccines in White-tailed deer (*Odocoileus virginianus*)

**DOI:** 10.1101/2024.09.20.614210

**Authors:** Julia Gonzalez, Cristina Harvey, Cárita de Souza Ribeiro-Silva, Brenda Leal-Galvan, Kelly A. Persinger, Pia U. Olafson, Tammi L. Johnson, Adela Oliva Chavez

## Abstract

Current tick control measures are focused on the use of synthetic acaricides and personal protective measures. However, the emergence of acaricide resistance and the maintenance of tick populations in wildlife has precluded the efficient management of ticks. Thus, host-targeted, non-chemical control measures are needed to reliably reduce ticks parasitizing sylvatic reservoirs. This project aimed to evaluate extracellular vesicles (EVs) from *Amblyomma americanum* as vaccine candidates for white-tailed deer (*Odocoileus virginianus*; WTD). Salivary gland (SG) and midgut (MG) EVs were isolated by ultracentrifugation. Three deer were vaccinated with SG and MG EVs and received two boosters at days 28 and 50. Two control deer were injected with adjuvant and PBS only. On day 58, WTD were infested with 100 *A. americanum* nymphs, 50 females, and 50 males that were allowed to feed to repletion. On-host and off-host mortality, tick engorgement weight, nymph molting, time to oviposition, and egg hatchability were evaluated. Serum samples were recovered every seven days until the last day of tick drop off, and then at one year (Y1) and 1-year and 1-month (Y1M1). Vaccination resulted in seroconversion and significant increases in total IgG levels that remained significantly higher than controls and pre-vaccination levels at Y1 and Y1M1. No negative effects were observed in nymphs, but on-host mortality of female *A. americanum* was significantly higher in vaccinated animals. No effects were observed on reproductive parameters. These results indicate that proteins within female tick SG and MG vesicles are not good candidates for vaccine design against nymphs; however, the on-host adult mortality suggests that tick EVs harbor protective antigens against *A. americanum* females.

## Introduction

Ticks, especially Ixodid (hard) ticks, continue to be a threat for animal and public health due to their ability to vector a great diversity of pathogens (Jongejan and Uilenberg, 2004). In the United States (US), the number of reported cases of tick-borne diseases is increasing each year (Paules et al., 2018). This increase, along with the expanding distributions of tick populations, heighten public health concerns (Eisen et al., 2017; Rochlin et al., 2022). The dynamics of wildlife host communities undoubtedly play an important role in the dispersal and/or the maintenance of ticks and, therefore, tick-borne pathogens. White-tailed deer (*Odocoileus virginianus*; WTD), particularly, are key in the life cycles of the lone star tick, *Amblyomma americanum,* and the black-legged tick, *Ixodes scapularis*, in the US (Tsao et al., 2021). They have different phenologies, but both species are three-host ticks that feed on a large diversity of animals and are vectors of pathogens that cause diseases of medical and veterinary importance, such as ehrlichiosis, rickettsiosis, Lyme disease, babesiosis and the illnesses caused by Heartland, Bourbon and Powassan viruses (Keirans et al., 1996; oddard and Varela-Stokes, 2009; Eisen and Eisen, 2018; Higuita et al., 2021). To control these tick populations, many efforts have focused on strategies to manage white-tailed deer, their main host, but there are still several issues associated with the management tools currently available (Stafford and Williams, 2017).

Tick control interventions target ticks on and/or off hosts and can involve ecological management, habitat modification, synthetic acaricide application, and immunological methods, such as anti-tick vaccines. However, all these measures have certain logistical limitations, including restrictions on areas to apply a treatment, potential detrimental effects on the environment and animal populations, and regulations on safe withdrawal time if game meat were to be consumed (White and Gaff, 2018). Although research on biological alternatives for tick control has been expanded in the last several decades (Quadros et al., 2020), chemical acaricides continue to be commonly applied. Their improper use can lead to negative ecological impacts and the emergence of acaricide resistance in tick populations. In fact, high tolerances to permethrin have been reported in *A. americanum* ticks recovered from farmed and wild deer in the US (Kaplan et al., 2022). Furthermore, the application of topical acaricides in wildlife, such as WTD, is difficult because they cannot be gathered as in the case of livestock. The treatment for WTD has been limited to administering an acaricide orally through food, that can be combined with a topical treatment, e.g. 4-poster device (Pound et al., 1996; Pound et al., 2000) or a motion-activated remote sprayer (Goolsby et al., 2022). Moreover, protocols to control the emergence and spread of chronic wasting disease (CWD) (Osterholm et al., 2019) has led to the prohibition of supplemental feeding of wild free-ranging deer in many US states, limiting the administration of oral acaricides and 4-post devices where deer may congregate. A different approach is excluding deer by fencing, which has also shown a significant reduction in *A. americanum* and *I. scapularis* tick populations (Bloemer et al., 1986; tafford, 1993; aniels and Fish, 1995). Elimination and reduction of WTD by culling remains questionable, as it has been somewhat effective in islands but was proven to be less effective in congested continental areas (Kugeler et al., 2016). Therefore, deer population reduction can be controversial and the control of tick populations using 4-poster devices and acaricides administered orally can be expensive, labor intensive, and is also not suitable for control in large areas (Wong et al., 2018; Nawrocki et al., 2023; Schulze et al., 2023). It is necessary to develop alternatives for tick management that are sustainable, economically feasible, and safe.

A promising alternative for tick control is the development of anti-tick vaccines. Current research efforts are focused on identifying tick antigens that could cause a host immune response, disturbing the tick attachment and/or feeding (Abbas et al., 2023). Following the advances in omics technology (transcriptomics, genomics, proteomics), several antigen-based vaccines have been developed; however, only a few have been tested in WTD. The commercially available antigen, Bm86, and Subolesin, have been tested as vaccine candidates against *Rhipicephalus microplus* in WTD (Carreón et al., 2012). Although these antigens resulted in a significant IgG conversion and decreased tick survival and fertility, Bm86 is ineffective against *Amblyomma cajennense* (Rodríguez-Valle et al., 2012) and Subolesin has only been tested in RNAi silencing experiments with *A. americanum*, which resulted in the lack of oviposition after microinjection of repleted females (Kocan et al., 2007). Furthermore, multivalent vaccines have been regarded as a potential improvement to anti-tick monovalent vaccines. However, the identification of synergistic antigens is important for this goal (Ndawula and Tabor, 2020). Synergistic antigens consist of proteins that, when injected together, increase tick control when compared to single antigen injection. Hence, it is essential to discover new antigens that can work together and act on multiple tick species to improve tick control.

Extracellular vesicles (EVs) are small lipid-rich particles secreted by eukaryotic cells and are known to participate in antigen transport and delivery (Sabanovic et al., 2021). Due to their biosafety and ability to enhance immunogenicity of proteins during vaccination, EVs are being explored as vaccine candidates against viruses (Sabanovic et al., 2021), parasites (Drurey et al., 2020), and cancer (Gonzalez-Melero et al., 2023). EVs were recently identified in tick saliva (Zhou et al., 2020) and are important in pathogen transmission and the modulation of the host skin immune system (Oliva Chávez et al., 2021; Butler et al., 2023), offering a new research field on the path to understanding tick-host interactions. For instance, it was reported recently that EVs from tick hemolymph, the tick’s blood, contain transport proteins potentially important in nutrient exchange between organs (Xu et al., 2023). Furthermore, proteomic analysis of tick salivary and hemolymph EVs (Nawaz et al., 2020; Oliva Chávez et al., 2021; Xu et al., 2023) identified several proteins previously tested as vaccine candidates packed within EVs, including Ferritin (Hajdusek et al., 2010) and glutathione S-transferase (Ndawula et al., 2019). Thus, we hypothesized that vaccination with tick EVs will result in strong humoral responses, inducing significant changes in antibody levels and controlling tick infestations on WTD. This study aims to evaluate the antigenicity of proteins found within EVs secreted by *A. americanum* tick salivary glands and midguts (*ex vivo* cultures) and examine the degree of tick control obtained by using these EVs to vaccinate WTD.

## Material and Methods

### Ethics statement

White-tailed deer experiments were performed at the Texas A&M AgriLife Research and Extension Center, Uvalde, TX, according to the Texas A&M AgriLife Institutional Animal Care and Use Committee (IACUC) approved protocol # 2021-029A (approved 11/16/2021).

### Tick infestation for EV production

*Amblyomma americanum* females were fed on three 1-year-old WTDs during three different infestations, as previously described (Baker et al., 2023). Briefly, the base of the neck, between the shoulder blades, was shaved and a stockinette was glued to the skin. One hundred *A. americanum* females were applied in the first two infestations and 150 females were applied in the last infestation. These infestations were separated from each other by approximately one month. Animals were housed individually to prevent allogrooming. Ticks were allowed to feed for 5 days and were removed with fine-tipped forceps. Salivary glands and midguts were removed/dissected and separately cultured ex vivo in vesicle-free tick media for 24 hours. EVs were isolated by serial centrifugation and followed by ultracentrifugation as described previously (Oliva Chávez et al., 2021) with a small modification. Supernatants were passed through a ReZist 1.0 µm syringe filter (Cytiva Life Sciences) prior to ultracentrifugation (Leal-Galvan et al., 2022). EVs from the first infestation were resuspended in 400 µl of PBS and aliquots were analyzed via Nanoparticle tracking analysis (NTA) and Transmission Electron Microscopy (TEM) as described below. A fourth tick infestation with 100 ticks/animal was performed in the same three WTD around one year after the first infestation, to produce more EVs needed for ELISAs.

### Nanoparticle tracking analysis (NTA)

An aliquot of 25 µl was taken from three replicates of vesicles secreted by the salivary glands or midguts dissected from 20 ticks. The 25 µl of vesicles were added to 475 µl of PBS for a 1:20 dilution. Vesicle concentrations and sizes were measured using a NanoSight LM10 (Malvern Panalytical) with NTA software version 3.2. We utilized a sCMOS camera and a Blue 405 laser; the camera level was set at 10 or 11 and the detect threshold was set at 5. Three reads of 60 minutes were performed. The average concentration for three technical replicates (three reads) and three biological replicates (three samples from 20 ticks) were visualized using GraphPad version 10.2.3 (Prism).

### Transmission Electron microscopy (TEM)

A second aliquot of 10 μl from three different samples was submitted to the Image Analysis Lab at Texas A&M School of Veterinary Medicine & Biomedical Sciences for the TEM visualization of the vesicles. Three replicates were visualized. Five microliters (5 μl) from each sample were pipetted on 300 Mesh, Formvar/Carbon Film, copper grids (Electron Microscopy Sciences) previously glow discharged for 30 seconds at 5 μA and incubated for 5 minutes. Excess liquid was blotted with filter paper and samples were rinsed twice with water. Vesicles were negatively stained with 200 µl of 1.5% phosphotungstic acid (PTA; Electron Microscopy Sciences). Excess liquid was blotted with filter paper and grids were air dried for at least 10 minutes. Visualization and imaging were done with a FEI Morgagni 268 transmission electron microscope (TEM) operated at 70kV, equipped with an ImageView III CCD camera. TEM images were collected using the iTEM software package. Image levels were adjusted with Adobe Photoshop.

### Vaccination

Vaccinations were done in three 1-year-old WTD (#924, #929 and #934) by injecting 400 μg of protein (200 μg from salivary gland EVs and 200 μg from midgut EVs) in 200 µl of PBS mixed with 200 µl of TiterMax Gold®, 1:1 ratio, intramuscularly. Two control deer (#923 and #930) were injected with TiterMax® Gold and PBS only (Figure 1). WTD used for the vaccination experiments had never been experimentally infested or vaccinated with tick antigens previously; nevertheless, whether they had been naturally exposed to ticks during captivity is unknown. All animals were boosted on days 28 and 50. Blood samples were collected on day 0 (pre-vaccination), every seven days post immunization and boost, at day 57 (pre-infestation), at the end of tick each tick infestation, which was dependent on the time to repletion (days 66-71 post-infestation), and then at one year (Y1) and one year and one month (Y1M1) post last booster (Figure 1). White-tailed deer blood samples were collected via jugular venipuncture using 10 ml red top vacuum blood collection tubes with 20-gauge collection needles.

**Figure 1.**
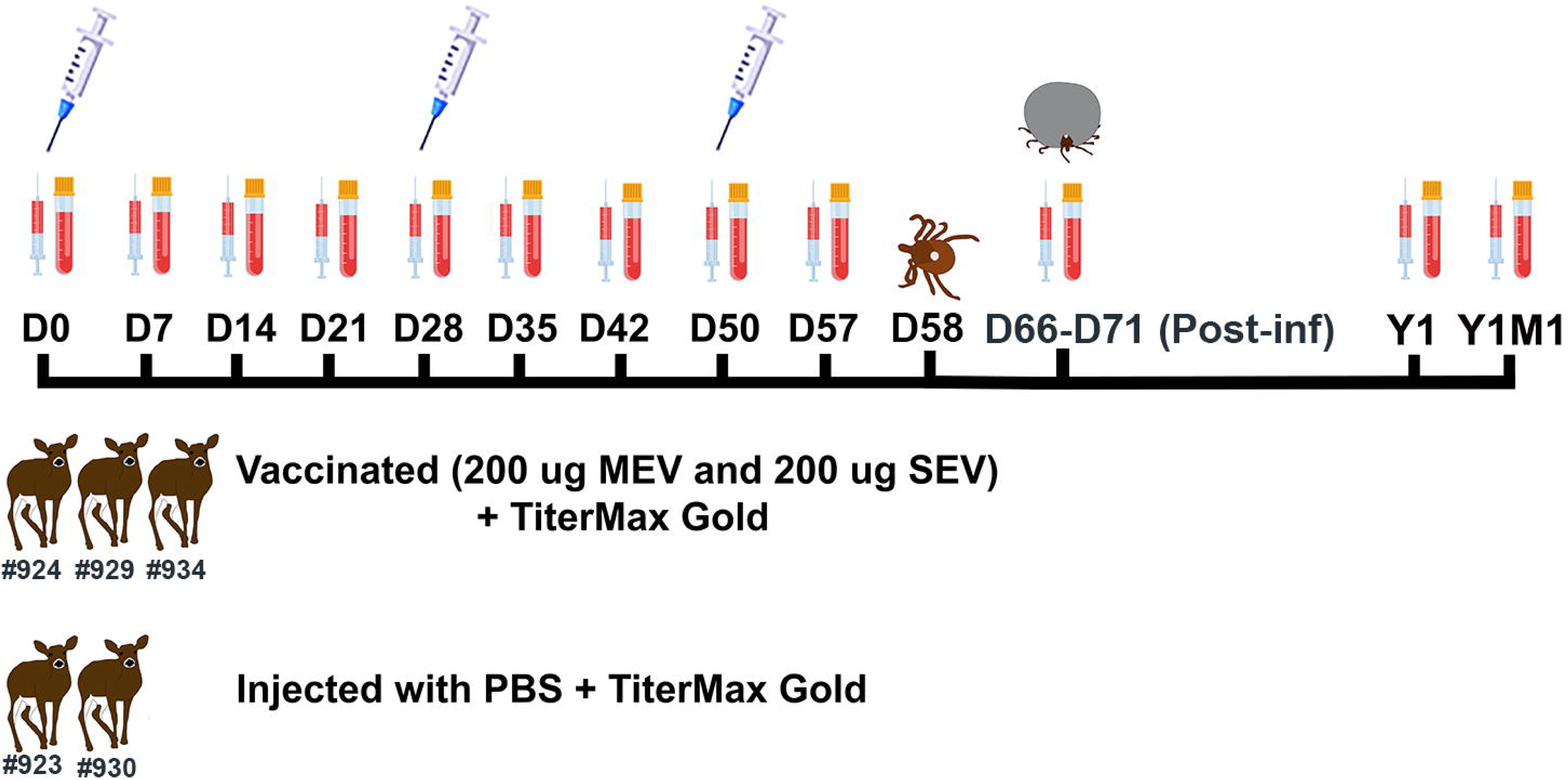
Experimental design and vaccination timeline. Three white-tailed deer were vaccinated with a combination of extracellular vesicles (EVs): 200 μg from midgut (MG) and 200 μg from salivary glands (SG), while two control animals received only PBS and adjuvant (TiterMax Gold). Blood samples were obtained at day 0 (pre-vaccination), every seven days post immunization and boost (syringe icon), pre-infestation (day 57), and post-infestation (day 69-70), one year (Y1) and one year and one month (Y1M1) post second booster. Ticks (100 *Amblyomma americanum* nymphs, 50 females, and 50 males) were applied on day 58 (8 days post second booster) and recovered after feeding to repletion.

### Tick infestation for vaccine evaluation

At 58 days after the first vaccination (Figure 1), WTD were infested with 100 *A. americanum* nymphs, 50 females, and 50 males provided from the pathogen free colony at Oklahoma State University and from USDA-ARS, Knipling-Bushland US Livestock Insects Research Laboratory (Kerrville, Texas). All ticks were allowed to feed to repletion. Nymphs were collected on day 5 and 6. *Amblyomma americanum* female collections occurred daily between days 8-13. For nymphs, 30 ticks were randomly selected from each group and weighted to determine differences in feeding. These nymphs were maintained in an incubator at 22 °C under 85% humidity in 16:8 h light/dark cycle to monitor molting. All recovered females that were alive were weighted and maintained for oviposition. Egg masses resulted were weighted and incubated at 25 °C under 85% humidity in 16:8 h light/dark cycle to allow hatching. Reproductive parameters were evaluated as previously described in (Baker et al., 2024). Pictures of adult ticks feeding on WTD were taken with a Super Speed Dual pixel 12 Megapixels F1.5 – 2.4 (dual aperture) camera. Auto tone in Adobe Photoshop was used in images from WTD #930 and WTD #934 day 10 to reduce redness.

We performed non-parametric Wilcoxon tests, or t-tests when possible, to compare the mean tick weight (tick engorgement) between the control and vaccinated group. Chi-square analyses were applied to evaluate significant differences between the number of ticks collected from each experimental group. Because of the small sample size included in this study (low expected frequencies with five white-tailed deer), we used Fisher’s exact tests to evaluate the significance of the association between tick categories (dead-feeding, dead-squished by deer, dead post-feeding, oviposited or molted and non-recovered) and the experimental groups (control and vaccinated); p-values were adjusted in this case by Holm-Bonferroni method to avoid the risk of Type I errors (false positives). All these analyses were performed using the software R Core Team (2021).

### Serum preparation

Whole blood was stored at room temperature for 4 to 6 hours, and then centrifuged at 2,000 xg for 10 minutes. The resultant serum was then aliquoted into cryovials and stored at-80 °C. Serum samples were shipped to College Station, TX or Madison, WI and stored at -80 °C until use.

### Antibody measurement by enzyme-linked immunosorbent assay (ELISA)

#### Sandwich ELISA

The total level of IgG in the serum was determined by sandwich ELISA. Rabbit anti-deer IgG (SeraCare) was diluted at 1:100 in ELISA coating buffer (BioLegend) and 100 µl added to each well in a 94-well Nunc™ MaxiSorp™ ELISA plate (BioLegend) and incubated overnight at 4 °C under agitation. Wells were washed three times with 200 µl of 1x ELISA wash buffer (BioLegend), and then 100 µl of 2% milk was added to each well for blocking. The plate was incubated for 1 h at room temperature. Serum samples were diluted 1:1,000 in 2% milk. Wells were washed three times with 200 µl of 1x ELISA wash buffer and 100 µl of diluted serum was added to each well. The plate was incubated with the diluted serum overnight at 4 °C under agitation. Wells were washed seven times with 200 µl of 1x ELISA wash buffer. Rabbit anti-deer IgG labeled with HRP (SeraCare) was diluted to 1:500 in 2% milk and 100 µl was added to each well. The plate was incubated overnight at 4 °C under agitation. Wells were washed ten times with 200 µl of 1x ELISA wash buffer and 100 µl TMB substrate kit (Thermo Scientific) was added to each well. The plate was incubated for 10 minutes at room temperature in a dark place. Development was stopped with 100 µl of ELISA stop solution (Invitrogen). Optical densities (ODs) were quantified with a NanoQuant Infinite M200 Pro (Tecan). No capture antibody, no serum, and no secondary were used as technical controls. Samples were run in duplicates. Statistical differences were evaluated on the OD averages from two independent experiments by Two-way ANOVA followed by Tukey test for multiple comparison in GraphPad version 10.2.3 (Prism).

To evaluate how long antibodies against EV proteins remain in circulation, antibodies in serum collected after Y1 and Y1M1 were measured and compared to day 0 (pre-vaccination), day 57 (pre-infestation), and post-infestation samples, using the same procedures as described above with the exception that wells were washed eight times with 200 µl of 1x ELISA wash buffer after the incubation with the secondary antibody and ODs were measured in a NanoQuant Infinite M1000 Pro (Tecan). No capture antibody, no serum, and no secondary were used as technical controls. Statistical differences were evaluated on the OD averages from two independent experiments by Two-way ANOVA followed by Tukey test for multiple comparison in GraphPad version 10.2.3 (Prism).

#### Indirect ELISA

The level of antibodies reactive to EV proteins were measured using an indirect ELISA. Salivary glands (SG) or midgut (MG) EVs were sonicated on ice for 5 seconds three times and then diluted 1:10 in PBS. Protein concentrations were measured using a Pierce™ BCA Protein Assay Kit (Thermofisher). Two hundred (200) ηg of protein per well were immobilized on a 94-well Nunc™ MaxiSorp™ ELISA plate overnight at 4 °C, using ELISA coating buffer (BioLegend). The rest of the procedure was performed as described above for the sandwich ELISA. The same procedure was used to measure long-lasting EV-specific antibodies, except that the plates were washed eight times with 200 µl of 1x ELISA wash buffer after the incubation with the secondary antibody and incubated for 15 minutes after the addition of TMB solution. No EVs, no serum, and no secondary were used as technical controls. Statistical differences were evaluated on the OD averages from two independent experiments by Two-way ANOVA followed by Tukey test for multiple comparison in GraphPad version 10.2.3 (Prism).

Differences in EV specific antibody titers at day 57 (pre-infestation) were compared using EV samples from salivary glands and midguts. Serum samples were diluted to 1:50, 1:100: 1:250, 1:500, 1:750, 1:1,000, 1:2,000, 1:4,000, 1:8,000, 1:10,000, 1:20,000, 1:50,000, and 1:100,000. No serum was used as technical control. Optical densities were measured in a NanoQuant Infinite M1000 Pro (Tecan). The titer average per animal from two plates was visualized as heat maps using GraphPad version 10.2.3 (Prism) except for titers against MG EVs for WTD #923, WTD #924, and WTD #934 that represent only one experimental replicate due to lack of sufficient vesicles to perform a second ELISA. The titer cut-off for all vaccinated sera samples was determined with the ODs of the control animals as negative control sera, as described (Frey et al., 1998), using the software R Core Team (2021).

## Results

### Salivary glands and midgut ex vivo cultures secrete a mixed population of extracellular vesicles

In mammalian systems, EVs are divided into exomeres (∼30 nm – 50 nm), small exosomes (60 nm – 80 nm), large exosomes (and 90 – 120 nm), and microvesicles (100 nm – 1 µm), depending on their biogenesis, cargo, and size distribution (Zhang et al., 2018). We recently developed a protocol for isolating of all these EV populations from tick *ex vivo* organ cultures (Leal-Galvan et al., 2022). In this study, we used NTA to confirm the size distribution of the vesicles secreted by each organ culture. Midgut EVs ranged from 51 nm – 550 nm with a peak size of 101 nm – 150 nm, corresponding to exosome and small microvesicle sizes (Figure 2A and Supplementary Video S1). Salivary gland EVs, on the other hand, had a size range from 51 nm – 500 nm with peaks at 101 nm – 150 nm and 151 nm – 200 nm in size, potentially representing exosomes and small microvesicles (Figure 2B and Supplementary Video S2). Transmission Electron Microscopy (TEM) visualization of midgut derived EVs confirmed the presence of vesicles of 50 nm – 200 nm, whereas TEM images of EVs secreted by salivary gland *ex vivo* cultures showed vesicles of 50 nm – 100 nm (Figure 2C). These results demonstrate that both organs predominantly secreted exosomes and microvesicles of small size, although exomeres were detected in low concentrations (Supplementary Files S1 and S2).

**Figure 2.**
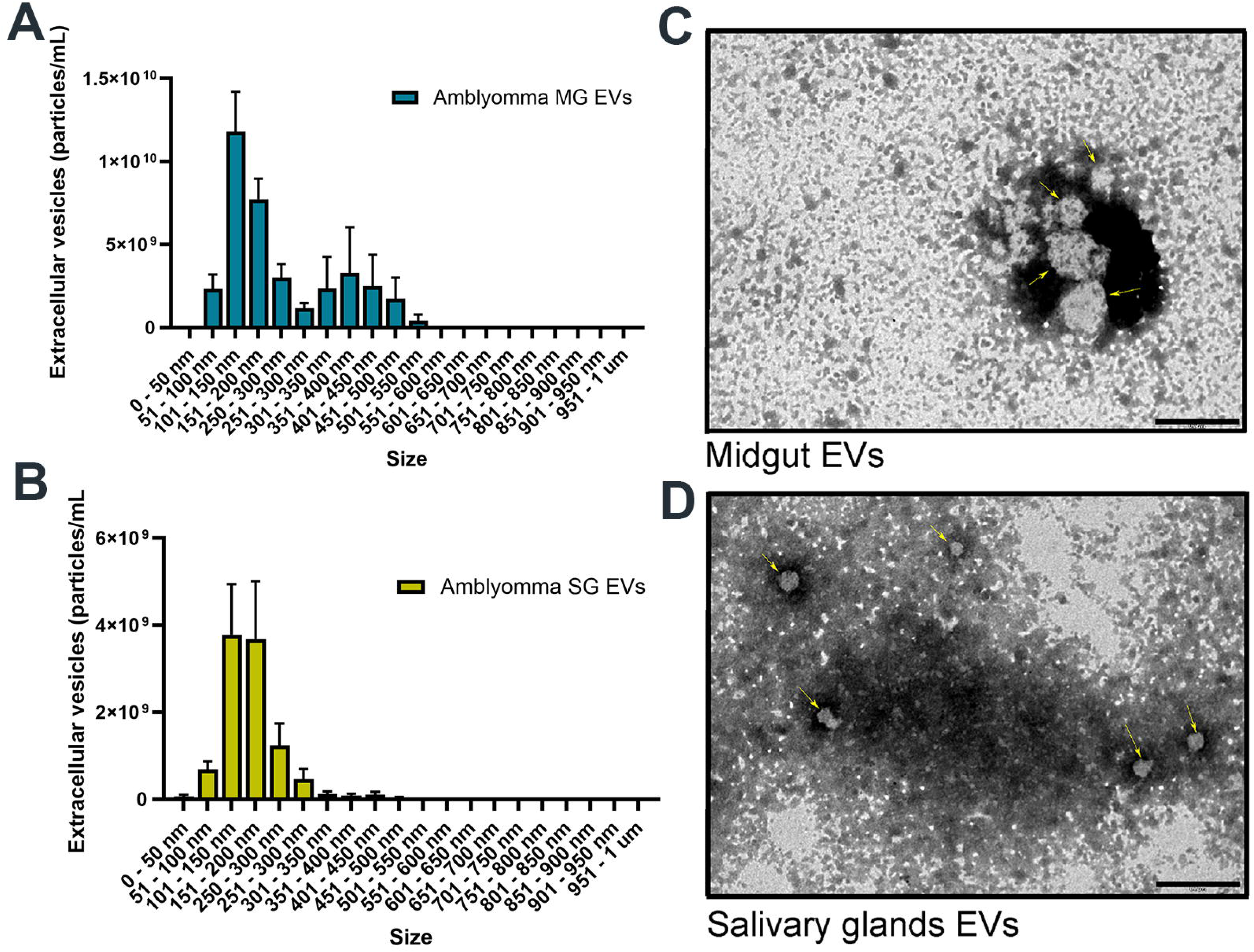
*Amblyomma americanum* midguts secreted a mixed population of extracellular vesicles, while *Amblyomma americanum* salivary glands secreted small vesicles. Nanotracking particle analysis (NTA) was used to measure the size and concentration of EVs secreted by *ex vivo* cultures of (A) midguts (MG) and (B) salivary glands (SG). Bars represent the average size of three technical replicates from three biological replicates (total of 9 measurements) ± the SEM. Transmission electron microscopy (TEM) was used to confirm size measurements. (C) Midgut EVs and (D) salivary glands EVs were visualized by negative staining with 1.5% phosphotungstic acid (PTA); yellow arrows point at vesicles and the size bar represents 0.2 µM.

### Vaccination with salivary and midgut extracellular vesicles results in seroconversion that is long-lasting

To determine the antigenic properties of tick EVs, we vaccinated WTD with *A. americanum* salivary and midgut EVs and measured circulating antibodies in the serum of vaccinated and control animals by sandwich ELISA. Total levels of deer IgG were not significantly different at any time point between vaccinated and control animals (p-value > 0.1; Supplementary Figure S1A and Supplementary File S3), except for WTD #929 that had significantly higher levels of total IgG when compared with control animal #923 at day 28 (p-value = 0.0167; Supplementary File S3). To measure EV-specific IgG antibodies, the serum from vaccinated and control animals were analyzed by indirect ELISAs. Vaccination resulted in a steady increase in anti-salivary and-midgut EV-specific antibodies with spikes after each boost (Figure 3A and B). Significant differences were detected in response to the vaccination to SG EVs (p-value < 0.0001; Supplementary File S4) and to MG EVs (p-value <0.0001; Supplementary File S5). Multiple comparison showed that anti-SG EV antibodies were significantly different in all vaccinated animals starting at day 35 and continued through post-infestation, when compared with both controls (p-value < 0.0011; Supplementary File S4). In the case of anti-MG EV antibodies, significant differences in antibody levels in all vaccinated animals, compared to the controls, were observed as early as day 14 and were still detectable until post-infestation (p-value < 0.0001; Supplementary File S5), suggesting that MG EVs might be more antigenic than SG EVs. Titer levels on day 57, which was seven days after the second boost and one day before tick infestation, were defined for each animal for SG EVs (Table 1; p-value ≤ 0.0005921; Supplementary File S6 and Figure S2A) and MG EVs (Table 2; p-value ≤ 2.925e-05; Supplementary File S7 and Figure S2B). WTD #924 and #934 had higher antibody titers against SG and MG EV proteins than WTD #929. All animals had higher antibody titers against MG EV proteins than SG EV proteins (Table 1 and 2), confirming that MG EV proteins are more antigenic than SG EV proteins.

**Figure 3.**
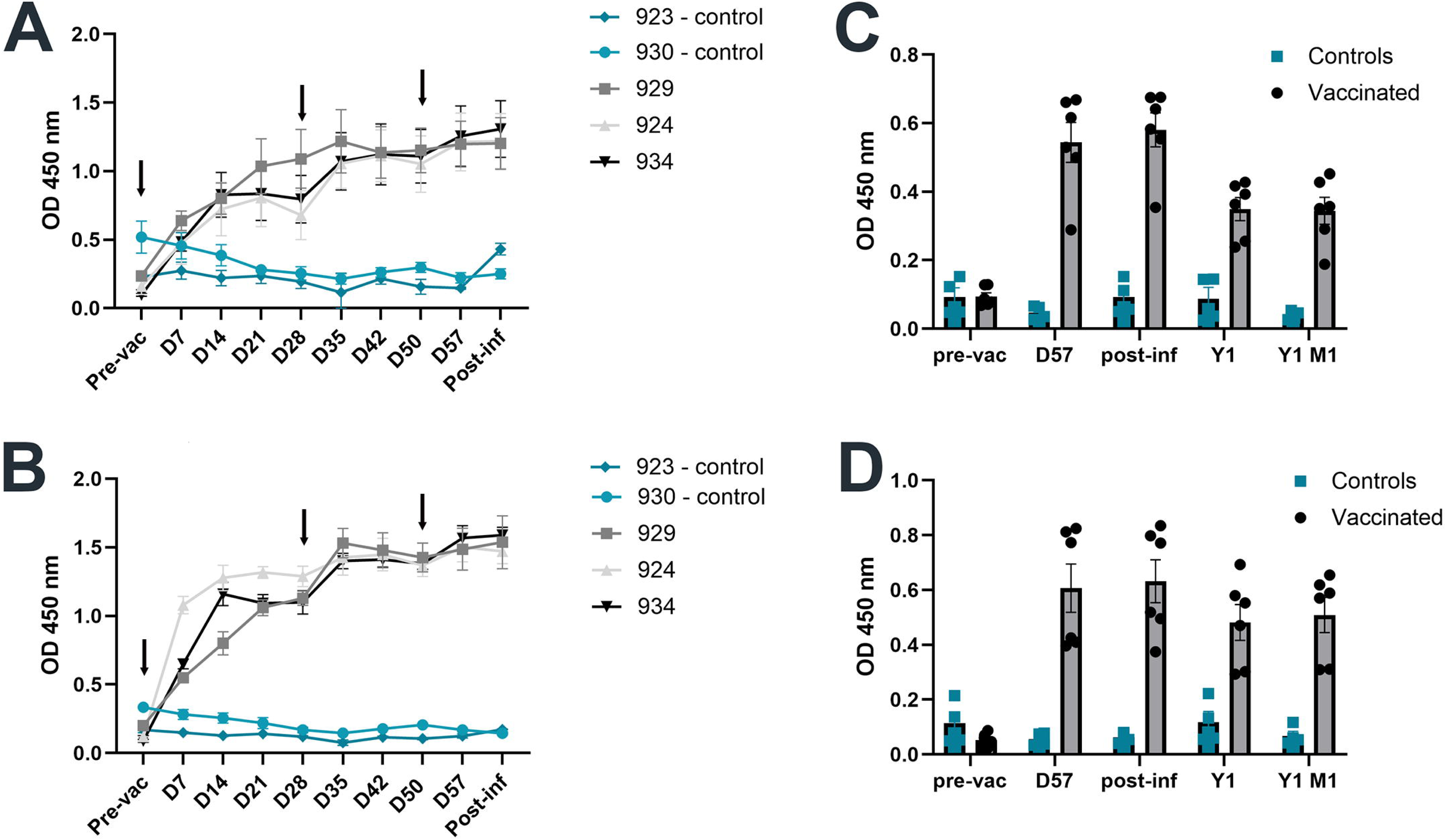
Vaccination results in increased and long lasting extracellular specific IgG levels. White-tailed deer were vaccinated with 200 µg of salivary EVs and 200 µg of midgut EVs. Serum samples were taken every 7 days to measure EV specific IgG levels by indirect ELISA, showing a steady response against (A) salivary EVs and (B) midgut EVs in all three vaccinated animals (black, light gray, and dark gray) versus control animals (light and dark blue). Lines represent the mean value of the average OD from two independent ELISAs ± SEM from individual animals. Statistical differences were evaluated by two-way ANOVA followed by Tukey multiple comparison. Black arrows indicate the days that injections were performed. Circulating antibody levels against (C) salivary and (D) midgut EVs were measured 1-year (Y1) and 1-year 1-month (Y1M1) later. Vaccinated animals (grey bars) showed higher antibody levels when compared to control animals (blue bars). Bars represent the average OD from two independent ELISAs ± SEM. The circles and squares represent the average reading from each animal in each ELISA. Statistical differences were evaluated by two-way ANOVA followed by Tukey multiple comparison.

**Table 1.** Titers of vaccinated animals against salivary glands EVs one day before tick infestation (D57)

| WTD # | Plate | Titer | Avg Titer | STDev |
| --- | --- | --- | --- | --- |
| 924 | 1 | 65832.59 | 46765.5 | 26965 |
|  | 2 | 27698.36 |  |  |
| 929 | 1 | 35769.7 | 24238.7 | 16307.3 |
|  | 2 | 12707.69 |  |  |
| 934 | 1 | 46776.97 | 38998.2 | 11000.8 |
|  | 2 | 31219.46 |  |  |

**Table 2.** Titers of vaccinated animals against midgut EVs one day before tick infestation (D57).

| WTD # | Plate | Titer | Avg Titer | STDev |
| --- | --- | --- | --- | --- |
| 924 | 1 | 100144.1 | N/A | N/A |
| 929 | 1 | 74191.72 | 74326.59 | 190.735 |
|  | 2 | 74461.46 |  |  |
| 934 | 1 | 77072.79 | N/A | N/A |

Previously commercialized anti-tick vaccines TickGARD^®^ and TickGARD^®*PLUS*^ were discontinued due to their short-lasting protection and the need for continuous boosts (3-4 per year) (abor, 2021). To assess whether EV derived antigens may lead to longer lasting effects, we measured the antibody levels one year (Y1) and one year and 1 month (Y1M1) after vaccination. Total levels of circulating IgG were not significantly different in any of the animals (p-value > 0.7781; Supplementary File S8 and Figure S1B). Nevertheless, significantly higher antibody levels were observed at Y1 and Y1M1 for both anti-SG EVs (Figure 3C; p-value < 0.0001) and anti-MG EVs (Figure 3D; p-value ≤ 0.0001) in vaccinated animals when compared to the controls and to each animal’s pre-vaccination levels (Supplementary Files S9 and S10). When considering individual animals, significantly higher levels of anti-SG EV antibodies were detected in all vaccinated animals in Y1 when compared to control WTD #923 (p-value ≤ 0.0015) and in WTD #924 versus control WTD #930 (p-value = 0.0071) and in all vaccinated animals versus all control animals in Y1M1 (p-value ≤ 0.0198). All vaccinated animals also showed significantly higher levels of circulating anti-SG EV antibodies at Y1 and Y1M1 when compared to pre-vaccinated levels (p-value ≤ 0.0392; Supplementary File S9 and igure S3A). On the other hand, only WTD #929 had significantly higher levels of antibodies against MG EV proteins in Y1 and Y1M1 when compared to control animals (p-value ≤ 0.018) and pre-vaccination antibody levels (p-value ≤ 0.0217; Supplementary File S10 and Figure S3B). Overall, these results demonstrate that EV vaccination leads to long lasting high levels of circulating antibodies, with some variation among individual animals.

### Vaccinations with female EVs does not affect nymph engorgement, but leads to on-host female mortality

To determine the effect of vaccination with MG and SG EVs, vaccinated and control WTD were infested with *A. americanum* nymphs and adults. Despite placing the ticks within sleeves for protection, some ticks were not collected due to the deer grooming a portion of the sleeves off and eating the ticks. These ticks were considered “non-recovered”, as they were not found on the animals or within the pens where the WTD were housed. The non-recovered ticks were not considered in further analyses because it was unknown whether they would fall within each of our final categories: engorged, unfed, dead during feeding (on-host) or post-feeding (off-host; dead prior to oviposition). One vaccinated animal (WTD #929) chewed off the patch containing nymphs and was eliminated from further nymphal analysis (Supplementary File S11). Most of the nymphs detached naturally after five days feeding, except 10 nymphs from a control WTD that were removed at 6 days post-infestation (dpi). We collected more dead nymphs from the control group than from the vaccinated group; however, these ticks were found detached from the deer, partially engorged and desiccated (personal observation). Similar numbers of nymphs were recovered from each group to evaluate the post-feeding effect (X-squared = 15.355, df = 1, p-value = 8.911e-05; Supplementary Figure S4A). There were no significant differences in the mean weight of a subsample of recovered nymphs (N = 30; control = 0.0077 ± 0.0004 g; vaccinated = 0.0080 ± 0.0003 g) (W = 1694.5, p-value = 0.5815; Supplementary Figure S4B). Interestingly, more nymphs molted from the vaccinated group than from the control group (X-squared = 13.955, df = 1, p-value = 0.0001873; Supplementary Files S11 and S12).

The *A. americanum* females started to detach from WTD at 8 dpi and ended on 13 dpi in both groups (Supplementary File S10 and Figure S5). One control animal (WTD #923) was excluded from the analysis because it removed the entire patch, losing a significant number of ticks (Supplementary File S11). We observed a similar pruritic response in the other animals, which resulted in collection of some squished females. The number of females dead during feeding, including those fatality squished, was significantly higher in the vaccinated group than the control group (adjusted p-value = 0.009250474) at 13 dpi (Figure 4A). This on-host mortality was observed in all three vaccinated animals. Some ticks were trapped by the hardening of exudate coming from the bite wound, especially those from WTD #929; ticks became stuck together, unable to feed, and had to be removed from the animal in scabbed clusters of dead ticks (Figure S6), which was not observed in the control animal. We started to collect dead-feeding females on 12 dpi, although in the case of WTD #934, some ticks were observed dead and visually desiccated, still attached to the animal, on 11 dpi (Figure 5). The mean weight of the females collected alive was similar in both groups (control =0.402±0.0472 g; vaccinated=0.360±0.0312 g) (W = 885, p-value = 0.376; Figure 4B).

**Figure 4.**
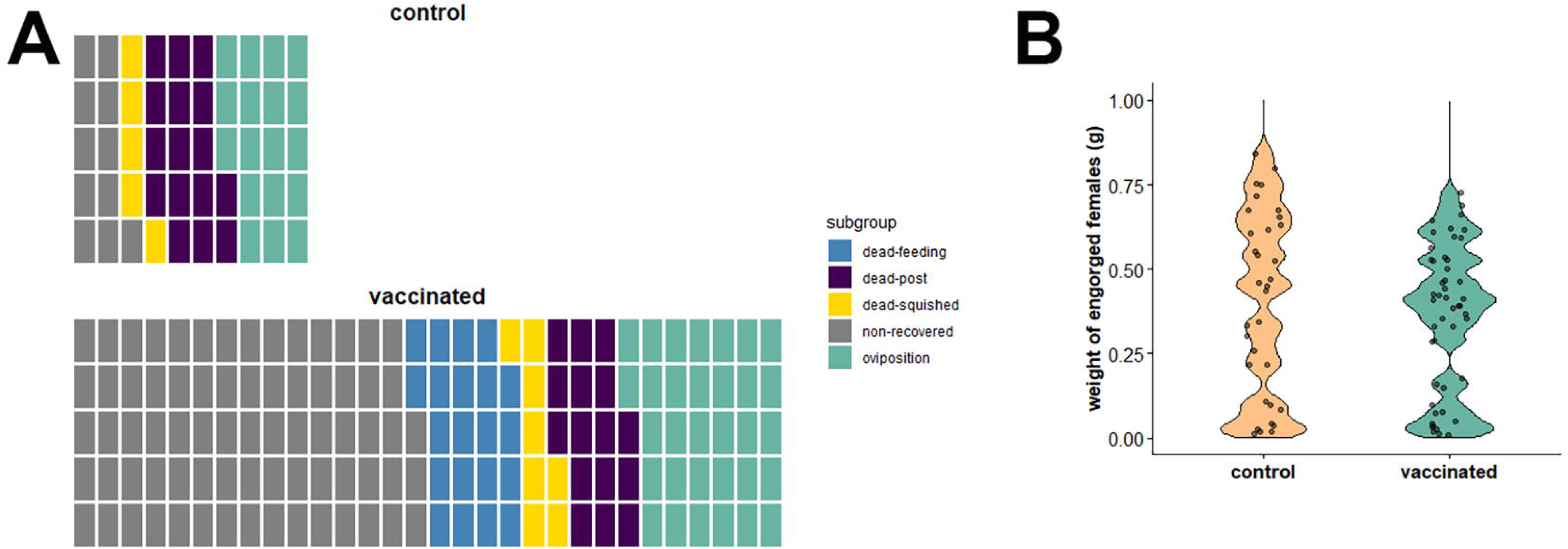
Vaccination with tick salivary and midgut extracellular vesicles leads to mortality during feeding but does not affect weight of surviving females. Tick infestation with *Amblyomma americanum* females on two groups of white-tailed deer: control (N=1) and vaccinated (N=3). Each white-tailed deer was infested with 50 females and 50 males. (A) We evaluated the number of females recovered during feeding, squished by the deer, post-feeding, and those that successfully started oviposition. Each square refers to one female. (B) Violin plots of the weights of female ticks collected from the control group (N=34 females) and vaccinated group (n=48 females).

**Figure 5.**
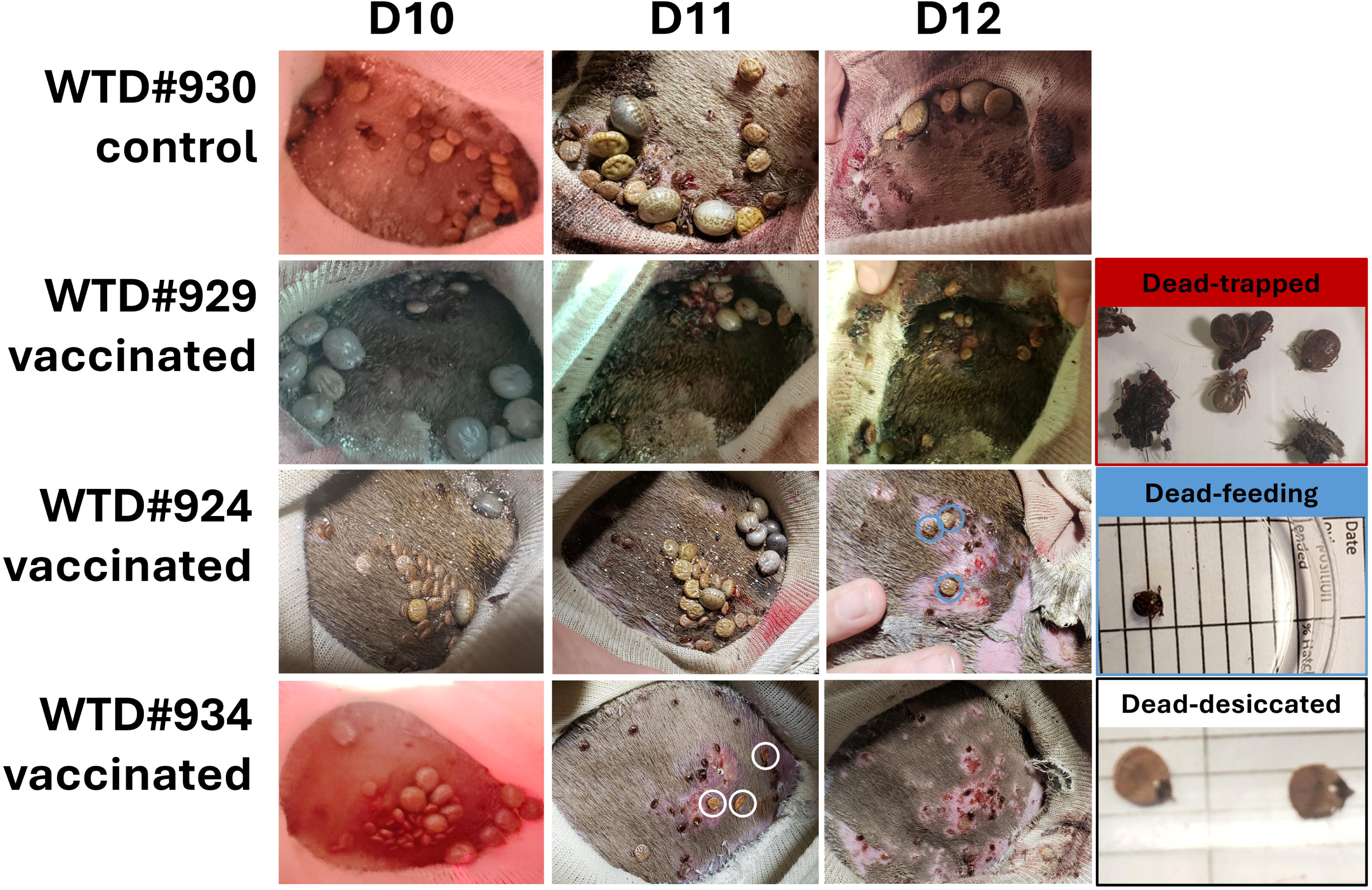
Pictorial progression of adult tick feeding in control and vaccinated animals. Pictures of the male and female *Amblyomma americanum* feeding on vaccinated and control animals were taken on day 10, 11, and 12 with a Super Speed Dual pixel 12 Megapixels F1.5 – 2.4 (dual aperture) camera. Several engorged females were observed in all patches. However, vaccinated animals (WTD #934, #929, and #924) presented ticks that died during feeding, some desiccated (white circles), apparently not feeding properly compared to other collected dead ticks (blue circles), or trapped dead due to hardened exudate from the bite site (red box).

To assess whether vaccination affected reproductive parameters of the ticks, the females that successfully fed and detached were maintained for up to two months to allow for oviposition (Supplementary File S11). The number of females that began oviposition was greater in the vaccinated group than in the control group, though females that died during oviposition were not significantly different between groups (Figure 4A; adjusted p-value = 0.253786825). Females that fed on vaccinated WTD #929 and #934 took longer to complete oviposition, however the differences between groups were not significant (control = 37.2±1.78 days; vaccinated = 41.1±1.33 days; t = - 1.8407, df = 33.678, p-value = 0.0745; Supplementary Files S11 and S12). Further, egg mass weight was not significantly different between groups (control = 0.234±0.0314 g; vaccinated = 0.230±0.0153 g; t = 0.11044, df = 25.254, p-value = 0.9129). We did not observe differences in the percentage of hatching (p-value = 0.5314; Figure 6A). A higher number of females produced offspring in the vaccinated group (N=29) than in the control group (N=17), yet there were not major differences in the average percentage of egg hatching (W = 274.5, p-value = 0.5314) or in the estimated number of larvae produced between groups (t = -0.11095, df = 26.676, p-value = 0.9125; control = 2444±423 larvae; vaccinated = 2498±244 larvae) (Figure 6B). Altogether our results indicate that vaccine effects are limited to on-host mortality and vaccination does not affect feeding, oviposition, or the number of larvae produced per female.

**Figure 6.**
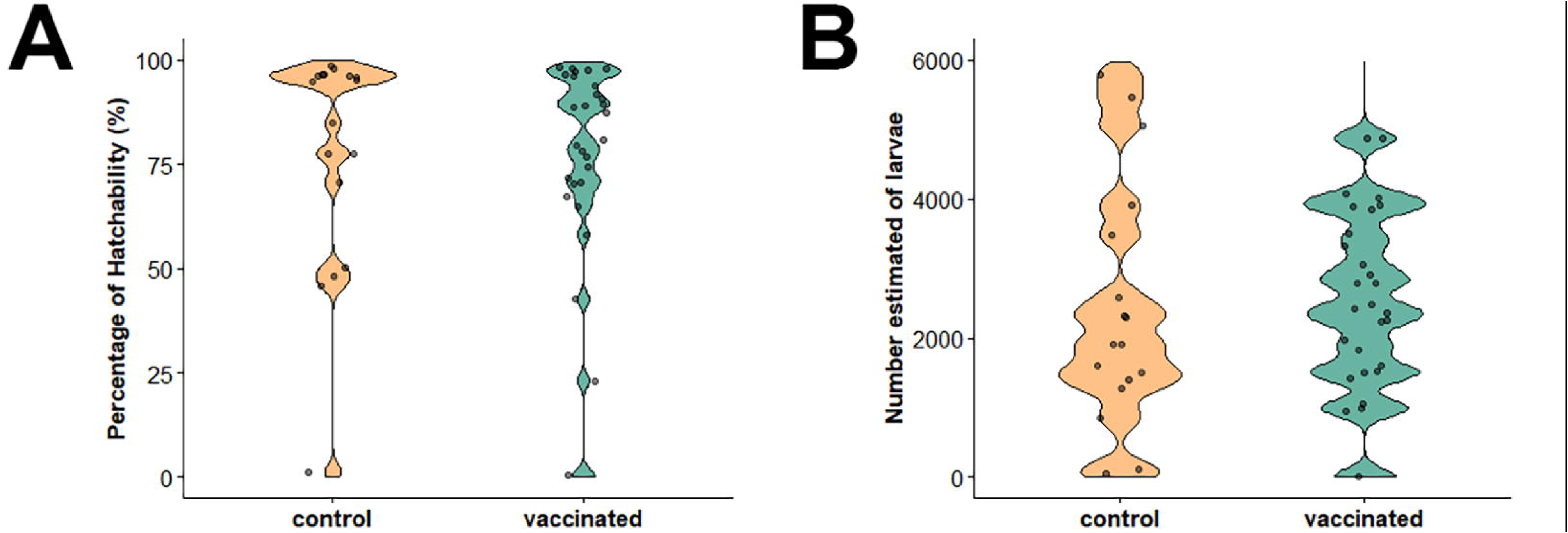
Vaccination with tick salivary and midgut extracellular vesicles does not affect *Amblyomma americanum* reproduction. Once the female completed the oviposition, it was removed, and egg masses were allowed to hatch for 60 days; at this time the contents were submerged in 70% ethanol. The dram was inverted 5 times and then spread onto a 10cm x 10 cm grid. Twenty-five cells were randomly selected, and the number of larvae and unhatched eggs was counted in each cell and multiplied by four to estimate the number of larvae hatched, number of eggs laid, and percent larval hatch. Violin plots of the egg development after oviposition from the control group (N=17 females) and vaccinated group (N=29 females). (A) Percentage of hatchability from the egg masses. (B) Estimated number of larvae hatched counted in 25 cells.

## Discussion

The role of exosomes as messengers within a cell-communication system makes them key players in important immunomodulatory processes such as, inflammation and activation of signaling pathways by carrying antigenic molecules, cytokines, and chemokines, and also by changing the metabolic reprograming of immune cells, serving a double purpose (Schorey and Harding, 2016; artins and Alves, 2020; Kalluri, 2024). Along with other studies, we previously showed that tick derived exosomes deliver immune-modulatory proteins that facilitate tick feeding, and these EVs are received by murine and human antigen presenting cells (APCs), such as macrophages (Nawaz et al., 2020; Oliva Chávez et al., 2021). Our current study demonstrates that *ex vivo* cultures of salivary glands and midguts from *A. americanum* females secrete a mixed population of EVs of small size, mainly exosomes and microvesicles (Figure 2). Remarkably, these EVs carry antigenic proteins that lead to long-lasting seroconversion in animals (Figures 3C and D). Exosome-based vaccines were first suggested in 1998 for cancer therapy (Zitvogel et al., 1998), but their use is now broadly applied in other research areas for the control of AIDS, hepatitis B, and other infectious and parasitic diseases (Santos and Almeida, 2021). The advantages of these vaccines include a diversity of antigens within the EV cargo or on their surface, presenting a wider spectrum of targets to stimulate several immune pathways. Parasite-derived EV vaccines also do not require attenuation, which can expedite safety trials (Sntos and Almeida, 2021). Although most of the studies related to immunization using parasite-derived EVs are still in the early stages (Alfandari et al., 2023), some studies have already shown promising results. This is the case of schistosomal EVs-based vaccine against schistosomiasis, that managed to significantly reduce the parasitic load and activated a specific antibody response in a mouse model (Mossallam et al., 2021).

In the case of ectoparasites, we reported recently that tick salivary EVs alter the frequency of dendritic epidermal T cells (DETCs), the cytokines and chemokines within murine skin (Oliva Chávez et al., 2021). However, further studies are needed to fully understand these mechanisms of action and develop vaccines to control ticks and tick-borne pathogen transmission. Some of the challenges faced by EV-based vaccine development are the large-scale production, the validation of their efficacy, and formulation, including investigating the dose required for safe and sufficient immune protection and selecting an adjuvant that leads to the development of protective immune responses (Stephenson et al., 2014; Santos and Almeida, 2021). In the present study, we used TiterMax® gold (TiterMax®, 2024), an oil/copolymer adjuvant that forms hydrophilic microparticules, which can increase the expression of Major Histocompatibility Complex II (MHC-II) and promote a balanced Th1/Th2 response. This mechanism leads to the production of IgG1 and IgG2a antibodies during vaccination with proteins from the endoparasite *Schistosoma mansoni* (Stephenson et al., 2014). Differences in resistance to tick infestations between cattle breeds are well known, including the potential association of specific MHC-II alleles (Acosta-Rodríguez et al., 2005; Untalan et al., 2007; Shyma et al., 2015). MHC-II, known as Bovine Leukocyte Antigen (BoLA) in cattle, is an antigen binding molecule that interacts with T cells, resulting in their activation and memory formation (Behl et al., 2012). Therefore, this is a key molecule in vaccination success. In fact, a deletion in BoLA-DRB3, an antigen recognition site that is highly polymorphic, increases the response of cattle to the previously commercially available TickGARD® (Sitte et al., 2002). The strong seroconversion observed in our results (Figures 3A and B; Figure S3) supports the utility of this adjuvant during anti-tick vaccination in WTD.

Extracellular vesicles from endoparasites contain antigenic proteins and vaccination can lead to the development of protective immunity (Shears et al., 2018). Ticks also secrete EVs into their saliva and hemolymph (Nawaz et al., 2020; Oliva Chávez et al., 2021; Xu et al., 2023), but their antigenic properties and applications are still under investigation. It is known that hard ticks (*Ixodidae*) display prolonged feeding, during which their saliva can suppress host immunity, facilitating feeding and pathogen transmission/acquisition (Francischetti et al., 2009; Kotál et al., 2015; Nuttall, 2023). Interestingly, some animal species can trigger an immune response against tick salivary components after repeated infestations, a phenomenon known as acquired tick resistance (ATR) that impacts tick biology, resulting in the detachment or even death of ticks (Kitsou et al., 2021; Narasimhan et al., 2021; van Oosterwijk and Wikel, 2021). Even though anti-tick resistance has not been reported in WTD populations, anti-tick immunity in natural hosts, such as cattle, has been observed during infestations with the winter tick, *Dermacentor albipictus*, a one-host tick that infests cattle and horses during the winter. Bishopp and Wood (1913) reported the resistance of a Jersey calf to *D. albipictus* that was evident by the formation of a scab that trapped the ticks at the bite site (Bishopp and Wood, 1913). Curiously, we observed the formation of such scabs in one of our animals (WTD #929) after vaccination with tick salivary and midgut EVs (Figure 5 and S6), which also presented the highest percentage of tick mortality during on-host feeding (Figure S5). Scab formation during tick infestation has been reported in Guinea pigs after multiple infestations. Histopathological examination of the bite site of these guinea pigs during the third infestation showed degranulating neutrophils and intense macrophage migration into the bite that led to a mix of macrophage-neutrophilic dermatitis (Anderson et al., 2017). We do not know what type of immune response led to the formation of the scabs in WTD #929 due to the lack of tools to study cellular immunity in deer; however, intense tissue damage was observed in the skin of the vaccinated animals (Figure 5). Our future studies will explore the cellular immune responses triggered in cattle, a model in which more tools are available, after vaccination with *A. americanum* SG and MG EVs. Nevertheless, our results highlight the need for more tools to study immune responses in wild-life reservoirs.

Interestingly, our results indicate that MG-EVs might be more antigenic than SG-EVs, evidenced by significantly different levels of anti-MG EV antibodies earlier than in anti-SG EV antibodies (Figures 3A and B). Moreover, the animals developed higher antibody titers to MG versus SG EVs (Table 1 and 2). Extracellular vesicles secreted by each organ type will likely carry tissue-specific proteins representing their cell of origin. Midgut antigens have long been studied for their ability to induce the production of antibodies (IgG) that interact with the midgut tissue, leading to its disruption. These antigens, termed as “concealed”, might avoid the immune-evasive nature of tick salivary secretions at the cost of requiring repeated vaccinations (Nuttall et al., 2006). In fact, we observed that although vaccination resulted in high levels of anti-MG-EV antibodies, only WTD #929 had significantly higher anti-MG EV IgG levels at Y1, whereas all animals showed significantly higher levels of anti-SG EV IgG levels Figure S2). Extracellular vesicles present in tick saliva during our experimental infestation might have served as a natural booster, leading to the maintenance of high IgG levels.

Vaccination with SG and MG EVs isolated from female *A. americanum* ticks led to significant differences in tick mortality during attachment (Figure 4A), but did not affect the engorgement of surviving ticks (Figure 4B) or their reproductive parameters (Figure 6). For this study, we did not weight ticks that died as we considered mortality as the end point. Likewise, we did not observe any significant effect on nymphal mortality or weight (Figure S4). It is unknown whether SG and MG EV content is different between life stages. A previous study showed significant differences in the protein content of saliva secreted by *Haemaphysalis longicornis* adults and nymphs. Only 31 proteins were shared between both life stages, with 30 proteins unique to nymphs and 74 proteins solely found in adults (Tirloni et al., 2015). The protein cargo within nymph and adult salivary EVs may be different enough that antibodies produced during vaccination with SG EVs from *A. americanum* females did not recognize nymph vesicles, but this hypothesis needs to be tested.

## Conclusion

Altogether, our study demonstrate that tick SG and MG EVs contain antigenic proteins that lead to significant seroconversion during vaccination, resulting in tick mortalities during attachment. Nevertheless, the use of these proteins for vaccination appears to only be effective against *A. americanum* females. Further research with EVs from nymphs is required for a vaccine development targeting this life stage. Our upcoming study will be focused on identifying specific antigenic proteins within EVs.

## Author contributions

AOC conceptualized the study; AOC and TLJ designed the experiments; AOC, KAP, and TLJ performed vaccinations; PUO provided ticks for vesicle production and evaluation of vaccine; KAP and TLJ performed tick infestations on WTD; AOC, JG, CSRS, CH, BLG, and TLJ dissected ticks for vesicle production; AOC assisted during the TEM imaging; JG, BLG, and AOC performed ELISAs of day 0 – post infection sera; AOC performed ELISAs on Y1/Y1M1 sera and titration ELISAs; AOC performed statistical analysis of ELISAs; JG performed evaluation of titers on R; AOC, KAP, and TLJ recovered nymphs and weighted; AOC randomly selected nymphs; KP and TLJ recovered adults and evaluated survival and reproduction; JG performed statistical analysis on tick parameters; JG, KAP, TLJ, and AOC wrote the manuscript; JG, AOC, TLJ, CSRS, BLG, and PUO edited the manuscript; all authors approved the final version of the manuscript.

## Acknowledgements

We thank Dr. Joseph Szule at the Image Analysis Lab in the College of Veterinary Medicine and Biomedical sciences for performing negative staining and TEM imaging and Charluz Rosario Arocho for performing NTA data acquisition. We thank Sydney Orsborn for helping to set up ticks during dissections and Sarah Durski and Stephanie Guzman Valencia for help during ELISAs at University of Wisconsin, Madison. Finally, we thank Teri Gaston and Alec Baker for their help with animal care and tick infestations.

## Abbreviations

EVs: Extracellular vesicles
MG: Midguts
SG: Salivary glands
US: United States
WTD: White-tailed deer
RNAi: RNA interference
pre-vac: Pre-vaccination
post-inf: Post-infestation
Y1: One year
Y1M1: One year and one month
ELISA: Enzyme-linked immunosorbent assay
ODs: Optical densities
Avg: Average
STDev: Standard deviation

---

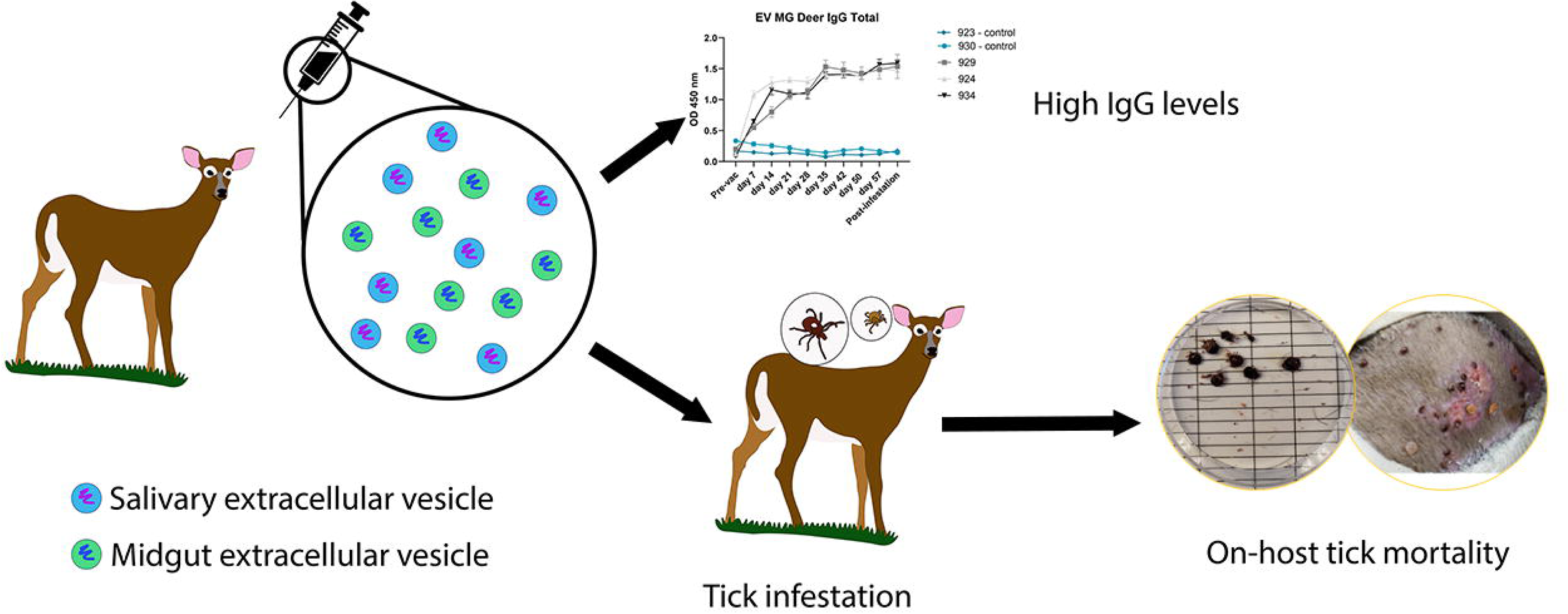

