## Supplementary figures and images for "Evaluation of tick salivary and midgut extracellular vesicles as anti-tick vaccines in White-tailed deer (*Odocoileus virginianus*)"

### Figure S1

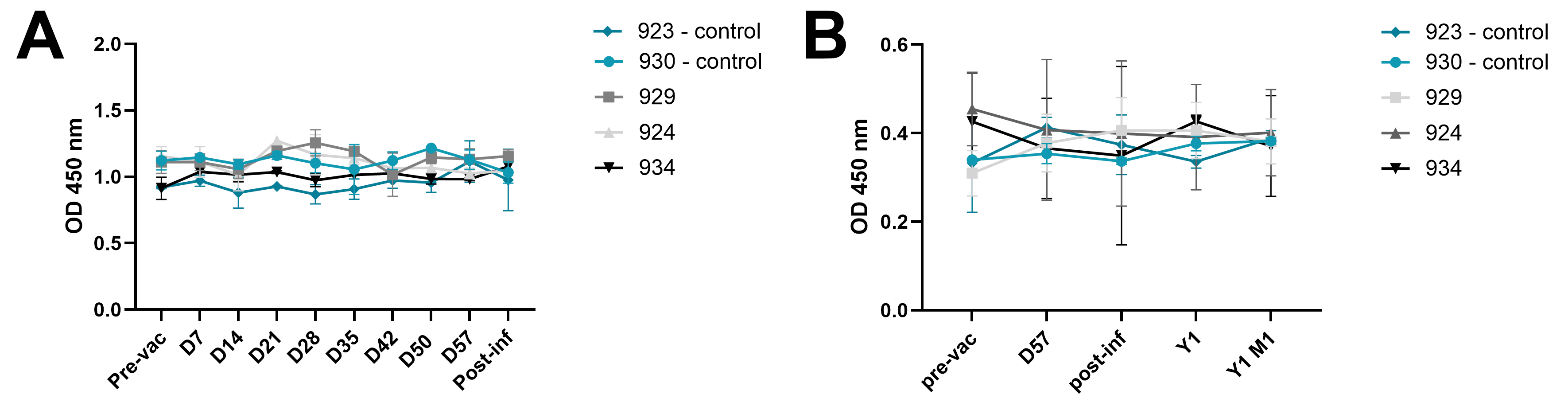

### Figure S2

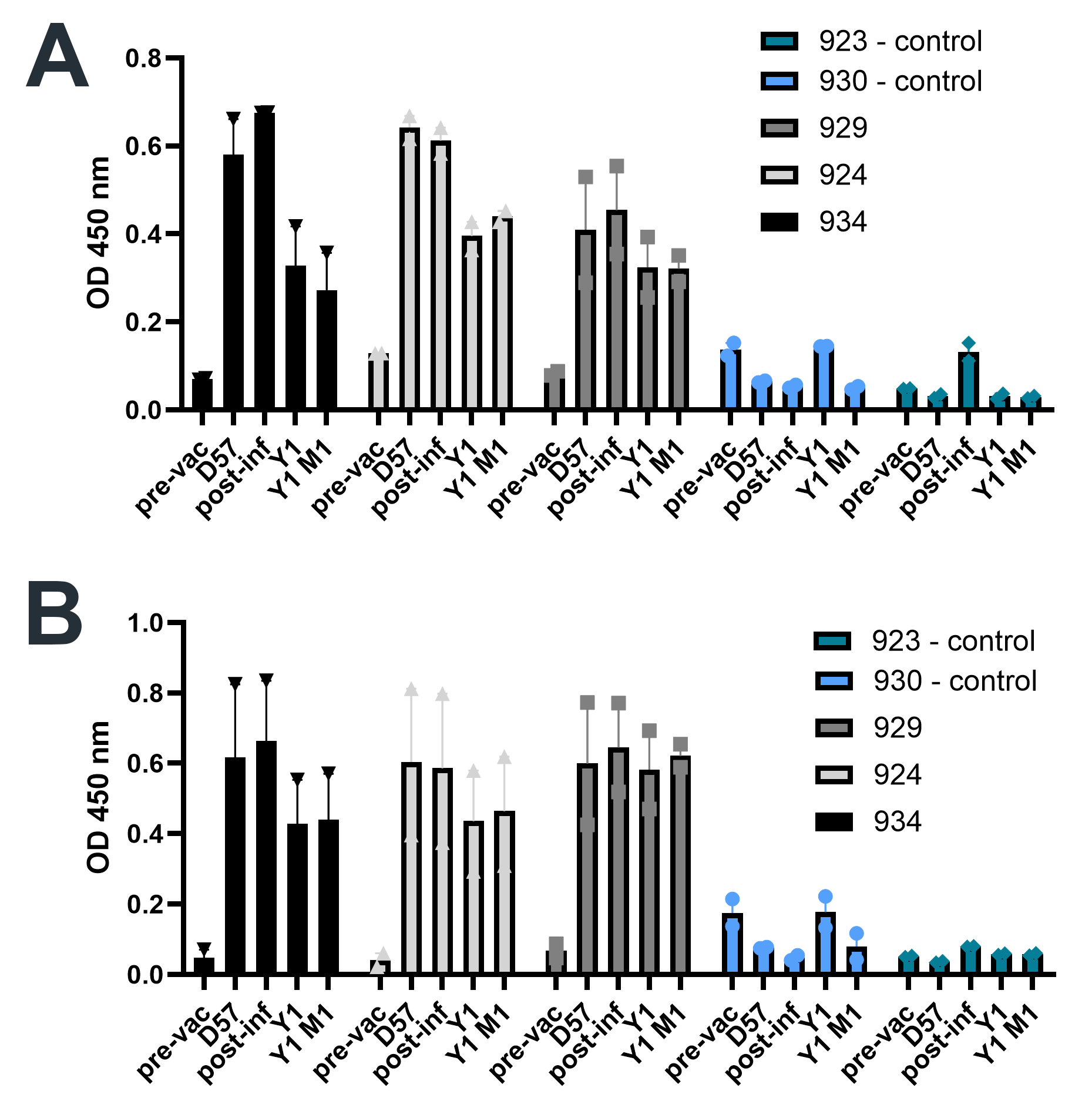

### Figure S3

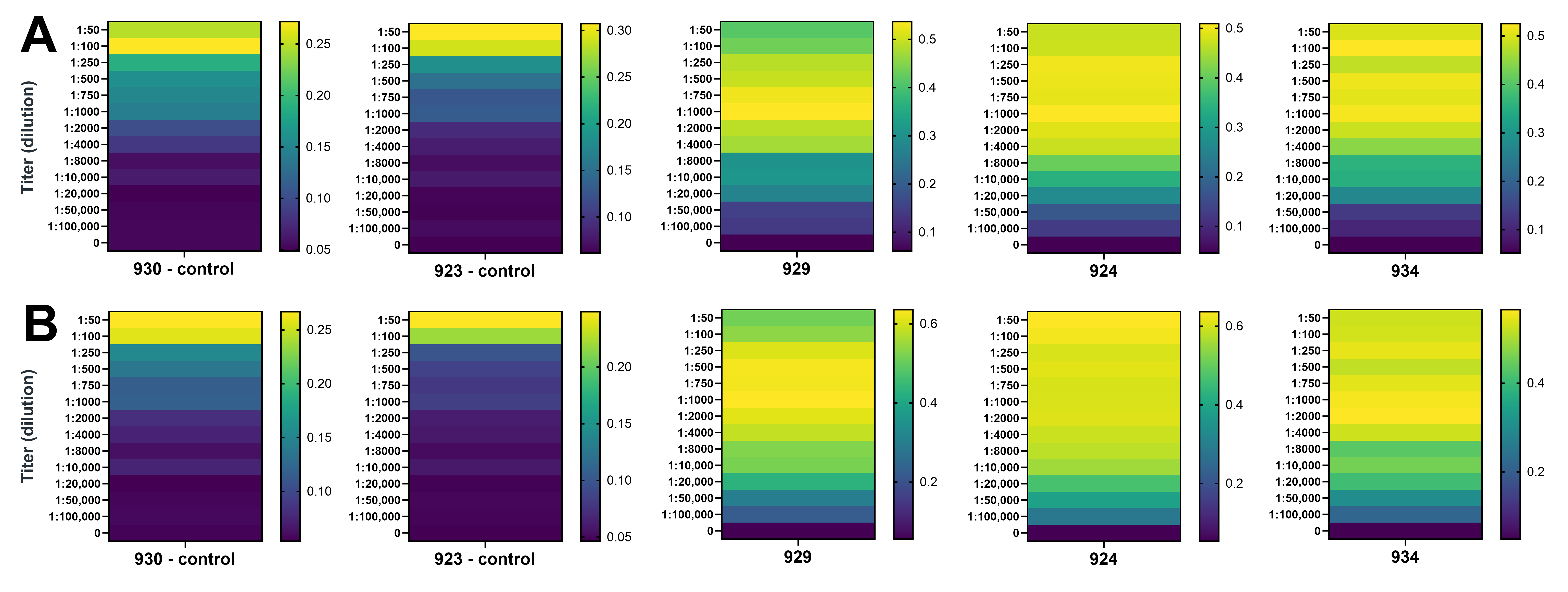

### Figure S4

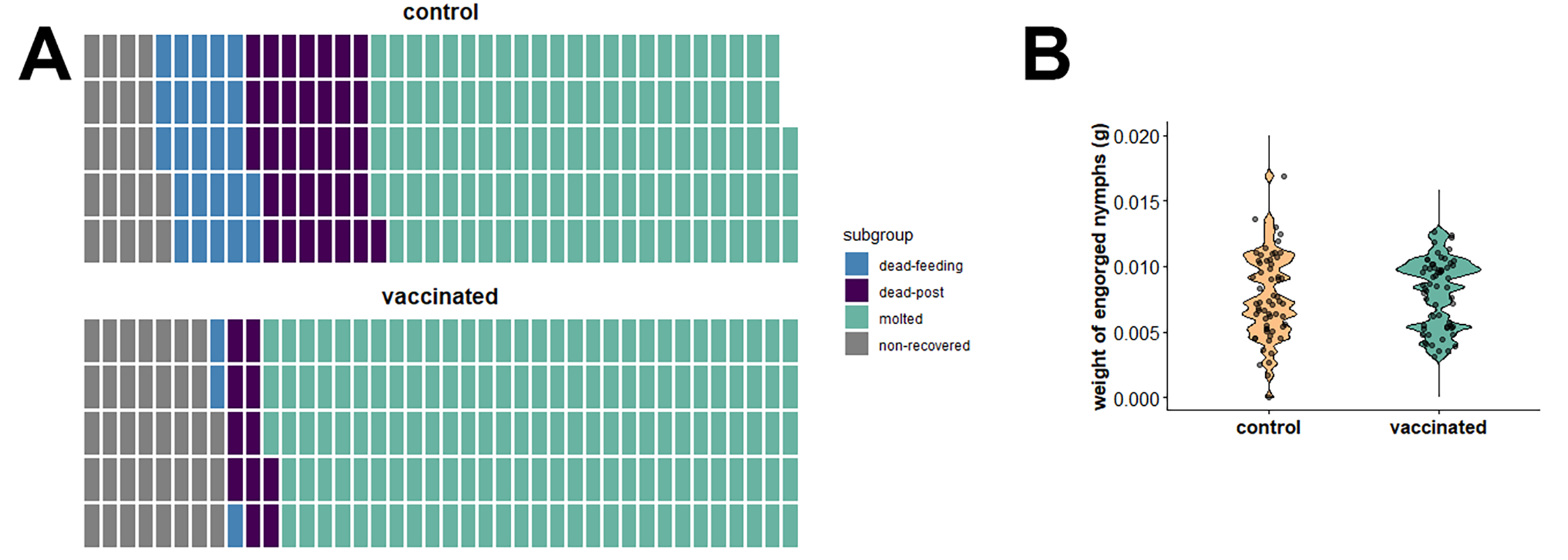

### Figure S5

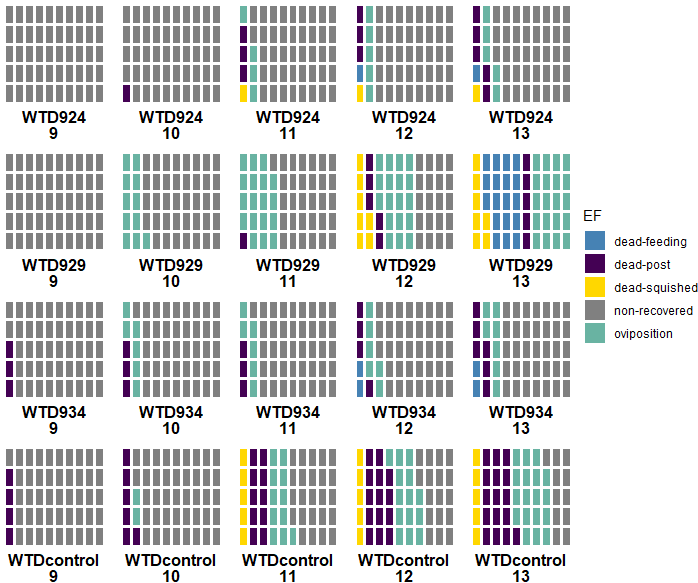

### Figure S6

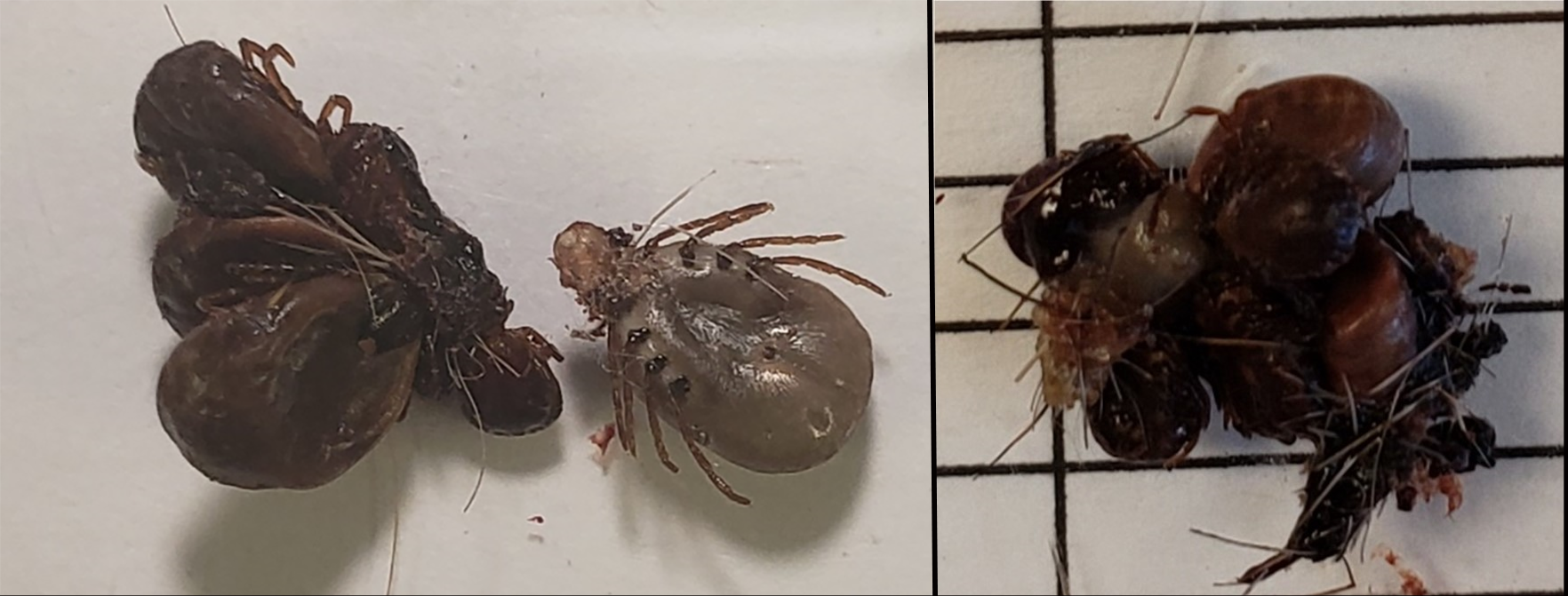
