## Supplemental figure legends for "Evaluation of tick salivary and midgut extracellular vesicles as anti-tick vaccines in White-tailed deer (*Odocoileus virginianus*)"

**Figure Captions**

**Figure S1. Total IgG levels in serum from white-tailed deer.** A sandwich ELISA was used to measure the total IgG levels circulating in control and vaccinated deer every (A) 7 days from pre-vaccination until post-infestation and (B) at one-year (Y1) and one-year and one-month (M1). Lines represent the mean value of the average OD from two independent ELISAs ± SEM from individual animals. Vaccinated animals are presented by light, dark and black lines, while control animals are shown in light and dark blue. Statistical differences were evaluated by two-way ANOVA followed by Tukey multiple comparison.

**Figure S2. Long-term individual levels of anti-EV IgGs in white-tailed deer.** Circulating antibody levels against (A) salivary and (B) midgut EVs were measured 1-year (Y1) and 1-year 1-month (Y1M1) later in each animal. Vaccinated animals (934 = black bars, 924 = light gray bars, and 929 = dark grey bars). Control animals (923 = dark blue bars and 930 = light blue bars). Bars represent the average OD from two independent ELISAs ± SEM. Statistical differences were evaluated by two-way ANOVA followed by Tukey multiple comparison.

**Figure S3. Heatmap representation of the IgG titers in white-tailed deer.** Serum samples from D57 (one day before infestation, were diluted at 1:50, 1:100: 1:250, 1:500, 1:750, 1:1,000, 1:2,000, 1:4,000, 1:8,000, 1:10,000, 1:20,000, 1:50,000, and 1:100,000. (A) Anti-SG EVs and (B) anti-MG EVs were measured by indirect ELISA. The average OD readings from two plates are represented as heatmaps, except for WTD 923, 924, and 934 that show the Average OD reading from 1 plate.

**Figure S4**. **Tick infestation with *Amblyomma americanum* nymphs on two groups of white-tailed deer**. control (N=2) and vaccinated (N=2). Each white-tailed deer was infested with 100 nymphs, except one control deer that was infested with 98 nymphs. (A) We evaluated the number of nymphs recovered during feeding, post-feeding, and the nymphs that successfully molted to adults. Each square refers to one nymph. (B) Violin plots of the weights of a subsample of nymphs collected from the control group (N=30) and vaccinated group (N=30).

**Figure S5**. **Tick infestation with *Amblyomma americanum* females on white-tailed deer**. control (N=1) and vaccinated (N=3). Each white-tailed deer was infested with 50 females and 50 males. We evaluated individually overtime the number of females recovered during feeding, squished by the deer, post-feeding, and those that successfully started oviposition since the first female detached (9 dpi). Each square refers to one female.

**Movie S1. Nanoparticle tracking analysis (NTA) video of midguts (MG) EVs.**

**Movie S2. Nanoparticle tracking analysis (NTA) video of salivary glands (SG) EVs.**

**Supplemental file S1. Nanoparticle tracking analysis (NTA) raw data of midguts (MG) EVs.**

**Supplemental file S2. Nanoparticle tracking analysis (NTA) raw data of salivary glands (SG) EVs.**

**Supplemental file S3. Raw data and statistical analysis of sandwich ELISA to measure total Deer IgG levels during vaccination.**

**Supplemental file S4. Raw data and statistical analysis of indirect ELISA to measure anti-SG EV Deer IgG levels during vaccination.**

**Supplemental file S5. Raw data and statistical analysis of indirect ELISA to measure anti-MG EV Deer IgG levels during vaccination.**

**Supplemental file S6. Titer calculation of anti-SG EV IgG in vaccinated white-tailed deer at day 57.**

**Supplemental file S7. Titer calculation of anti-MG EV IgG in vaccinated white-tailed deer at day 57.**

**Supplemental file S8. Raw data and statistical analysis of sandwich ELISA to measure total Deer IgG levels at one year and one year and one month post vaccination.**

**Supplemental file S9. Raw data and statistical analysis of indirect ELISA to measure anti-SG EVs IgG levels at one year and one year and one month post vaccination.**

**Supplemental file S10. Raw data and statistical analysis of indirect ELISA to measure anti-MG EVs IgG levels at one year and one year and one month post vaccination.**

**Supplemental file S11. Raw data tick infestation on white-tailed deer.**

**Supplemental file S12. Statistical analysis of results from tick infestation on white-tailed deer.**
