## Supplemental file S6 for "Evaluation of tick salivary and midgut extracellular vesicles as anti-tick vaccines in White-tailed deer (*Odocoileus virginianus*)"

### cutoff-and-titer_ELISA.R

gonzalezj 2024-07-03

| WTD | Titer |
|---|---|
| 924 – P1 | 65832.59 |
| 924-P2 | 27698.36 |
| 929 -P1 | 35769.7 |
| 929 – P2 | 12707.69 |
| 934 -P1 | 46776.97 |
| 934 -P2 | 31219.46 |

```
##ELISA titer determination
#924-P1
> cutoff= 0.2608443
> OD=read.table("clipboard")
> OD
       V1
1  0.51815
2  0.51100
3  0.52285
4  0.52060
5  0.51455
6  0.54565
7  0.52060
8  0.51090
9  0.41960
10 0.45385
11 0.37755
12 0.29235
13 0.20440
> Dilutions=c(50,100,250,500,750,1000,2000,4000,8000,10000,20000,50000,100000)
>
> #Different approaches to get significant p-value and Rsquare >0.7
> Regression=lm(Dilutions~OD$V1)
> log10_Regression=lm((log10(Dilutions))~OD$V1)
> Reciprocal_Regression=lm((1/Dilutions)~OD$V1)
>
> summary(Regression)

Call:
lm(formula = Dilutions ~ OD$V1)

Residuals:
   Min     1Q Median     3Q    Max 
-16324  -5368   1494   3548  19409 

Coefficients:
            Estimate Std. Error t value Pr(>|t|)    
(Intercept)   134034      12866  10.418 4.90e-07 ***
OD$V1        -261465      27622  -9.466 1.28e-06 ***
---
Signif. codes:  0 ‘***’ 0.001 ‘**’ 0.01 ‘*’ 0.05 ‘.’ 0.1 ‘ ’ 1

Residual standard error: 10030 on 11 degrees of freedom
Multiple R-squared:  0.8907,	Adjusted R-squared:  0.8807 
F-statistic: 89.61 on 1 and 11 DF,  p-value: 1.276e-06
```

```
> Titre_1=coef(Regression)[1] + (cutoff)*coef(Regression)[2]
> Titre_1
(Intercept)
   65832.59
>
> summary(log10_Regression)

Call:
lm(formula = (log10(Dilutions)) ~ OD$V1)

Residuals:
     Min       1Q   Median       3Q      Max
-1.13391 -0.36516  0.04365  0.38908  0.71067

Coefficients:
            Estimate Std. Error t value Pr(>|t|)
(Intercept)   7.0149     0.7628   9.196  1.7e-06 ***
OD$V1        -8.0710     1.6377  -4.928 0.000451 ***
---
Signif. codes:  0 '***' 0.001 '**' 0.01 '*' 0.05 '.' 0.1 ' ' 1

Residual standard error: 0.5944 on 11 degrees of freedom
Multiple R-squared:  0.6883,	Adjusted R-squared:   0.66
F-statistic: 24.29 on 1 and 11 DF,  p-value: 0.0004508

> A=coef(log10_Regression)[1] + (cutoff)*coef(log10_Regression)[2]
> Titre_2=10^A
> Titre_2
(Intercept)
   81208.15
>
> summary(Reciprocal_Regression)

Call:
lm(formula = (1/Dilutions) ~ OD$V1)

Residuals:
       Min         1Q     Median         3Q        Max
-0.0038183 -0.0029128 -0.0022478 -0.0000048  0.0157976

Coefficients:
             Estimate Std. Error t value Pr(>|t|)
(Intercept) -0.005384   0.007328  -0.735    0.478
OD$V1        0.018501   0.015733   1.176    0.264

Residual standard error: 0.00571 on 11 degrees of freedom
Multiple R-squared:  0.1117,	Adjusted R-squared:  0.03092
F-statistic: 1.383 on 1 and 11 DF,  p-value: 0.2644

> B=coef(Reciprocal_Regression)[1] + (cutoff)*coef(Reciprocal_Regression)[2]
> Titre_3=1/B
> Titre_3
(Intercept)
  -1791.875
```

```
#924-P2
> OD=read.table("clipboard")
> OD
        V1
1  0.42305
2  0.43710
3  0.47610
4  0.47300
5  0.47045
6  0.47320
7  0.45305
8  0.43070
9  0.38490
10 0.22540
11 0.15145
12 0.04960
13 0.04885
> Dilutions=c(50,100,250,500,750,1000,2000,4000,8000,10000,20000,50000,100000)
>
> #Different approaches to get significant p-value and Rsquare >0.7
> Regression=lm(Dilutions~OD$V1)
> log10_Regression=lm((log10(Dilutions))~OD$V1)
> Reciprocal_Regression=lm((1/Dilutions)~OD$V1)
>
> summary(Regression)

Call:
lm(formula = Dilutions ~ OD$V1)

Residuals:
   Min     1Q Median     3Q    Max
-23865  -3677   1403   4155  40973

Coefficients:
            Estimate Std. Error t value Pr(>|t|)
(Intercept)    66247      10899   6.078 7.98e-05 ***
OD$V1        -147782      28636  -5.161 0.000313 ***
---
Signif. codes:  0 '***' 0.001 '**' 0.01 '*' 0.05 '.' 0.1 ' ' 1

Residual standard error: 16390 on 11 degrees of freedom
Multiple R-squared:  0.7077,  Adjusted R-squared:  0.6811
F-statistic: 26.63 on 1 and 11 DF,  p-value: 0.000313

> Titre_1=coef(Regression)[1] + (cutoff)*coef(Regression)[2]
> Titre_1
(Intercept)
   27698.36
>
> summary(log10_Regression)

Call:
lm(formula = (log10(Dilutions)) ~ OD$V1)

Residuals:
```

```
     Min       1Q   Median       3Q      Max 
-1.25078 -0.16139  0.03906  0.30683  0.75817 

Coefficients:
            Estimate Std. Error t value Pr(>|t|)    
(Intercept)   5.1141     0.3955  12.932 5.37e-08 ***
OD$V1        -5.1161     1.0391  -4.924 0.000454 ***
---
Signif. codes:  0 '***' 0.001 '**' 0.01 '*' 0.05 '.' 0.1 ' ' 1

Residual standard error: 0.5948 on 11 degrees of freedom
Multiple R-squared:  0.6879,	Adjusted R-squared:  0.6595 
F-statistic: 24.24 on 1 and 11 DF,  p-value: 0.0004541

> A=coef(log10_Regression)[1] + (cutoff)*coef(log10_Regression)[2]
> Titre_2=10^A
> Titre_2
(Intercept) 
   6020.147 
> 
> summary(Reciprocal_Regression)

Call:
lm(formula = (1/Dilutions) ~ OD$V1)

Residuals:
       Min         1Q     Median         3Q        Max 
-0.0037042 -0.0033299 -0.0016161  0.0002186  0.0161292 

Coefficients:
              Estimate Std. Error t value Pr(>|t|)
(Intercept) -0.0007411  0.0038293  -0.194    0.850
OD$V1        0.0109015  0.0100612   1.084    0.302

Residual standard error: 0.005759 on 11 degrees of freedom
Multiple R-squared:  0.09644,	Adjusted R-squared:  0.0143 
F-statistic: 1.174 on 1 and 11 DF,  p-value: 0.3018

> B=coef(Reciprocal_Regression)[1] + (cutoff)*coef(Reciprocal_Regression)[2]
> Titre_3=1/B
> Titre_3
(Intercept) 
   475.6223 

#929-P1
> OD=read.table("clipboard")
> OD
        V1
1  0.34870
2  0.35655
3  0.43205
4  0.39900
5  0.47100
6  0.47855
7  0.45655
```

```
8  0.43090
9  0.34250
10 0.35060
11 0.28535
12 0.12705
13 0.15015
> Dilutions=c(50,100,250,500,750,1000,2000,4000,8000,10000,20000,50000,100000)
>
> #Different approaches to get significant p-value and Rsquare >0.7
> Regression=lm(Dilutions~OD$V1)
> log10_Regression=lm((log10(Dilutions))~OD$V1)
> Reciprocal_Regression=lm((1/Dilutions)~OD$V1)
>
> summary(Regression)

Call:
lm(formula = Dilutions ~ OD$V1)

Residuals:
   Min     1Q Median     3Q    Max
-16675 -10458  -5322   8653  40235

Coefficients:
            Estimate Std. Error t value Pr(>|t|)
(Intercept)    92313      15742   5.864 0.000109 ***
OD$V1        -216770      42305  -5.124 0.000331 ***
---
Signif. codes:  0 ‘***’ 0.001 ‘**’ 0.01 ‘*’ 0.05 ‘.’ 0.1 ‘ ’ 1

Residual standard error: 16470 on 11 degrees of freedom
Multiple R-squared:  0.7047,  Adjusted R-squared:  0.6779
F-statistic: 26.25 on 1 and 11 DF,  p-value: 0.0003314

> Titre_1=coef(Regression)[1] + (cutoff)*coef(Regression)[2]
> Titre_1
(Intercept)
    35769.7
>
> summary(log10_Regression)

Call:
lm(formula = (log10(Dilutions)) ~ OD$V1)

Residuals:
    Min      1Q  Median      3Q     Max
-1.6891 -0.3908  0.3819  0.5372  0.7014

Coefficients:
            Estimate Std. Error t value Pr(>|t|)
(Intercept)   5.4560     0.7695   7.090 2.02e-05 ***
OD$V1        -5.9303     2.0681  -2.868   0.0153 *
---
Signif. codes:  0 ‘***’ 0.001 ‘**’ 0.01 ‘*’ 0.05 ‘.’ 0.1 ‘ ’ 1

Residual standard error: 0.8053 on 11 degrees of freedom
```

```
Multiple R-squared:  0.4278,  Adjusted R-squared:  0.3758
F-statistic: 8.223 on 1 and 11 DF,  p-value: 0.01531

> A=coef(log10_Regression)[1] + (cutoff)*coef(log10_Regression)[2]
> Titre_2=10^A
> Titre_2
(Intercept)
   8112.063
>
> summary(Reciprocal_Regression)

Call:
lm(formula = (1/Dilutions) ~ OD$V1)

Residuals:
      Min        1Q    Median        3Q       Max
-0.003066 -0.002853 -0.002233 -0.001194  0.016998

Coefficients:
            Estimate Std. Error t value Pr(>|t|)
(Intercept) 0.001670   0.005773   0.289    0.778
OD$V1       0.003820   0.015516   0.246    0.810

Residual standard error: 0.006042 on 11 degrees of freedom
Multiple R-squared:  0.005481,      Adjusted R-squared:  -0.08493
F-statistic: 0.06062 on 1 and 11 DF,  p-value: 0.8101

> B=coef(Reciprocal_Regression)[1] + (cutoff)*coef(Reciprocal_Regression)[2]
> Titre_3=1/B
> Titre_3
(Intercept)
    375.082

#929-P1
> OD=read.table("clipboard")
> OD
        V1
1  0.47665
2  0.51005
3  0.54230
4  0.59325
5  0.58170
6  0.59495
7  0.52095
8  0.51415
9  0.26175
10 0.26985
11 0.25865
12 0.17155
13 0.12490
> Dilutions=c(50,100,250,500,750,1000,2000,4000,8000,10000,20000,50000,100000)
>
> #Different approaches to get significant p-value and Rsquare >0.7
> Regression=lm(Dilutions~OD$V1)
> log10_Regression=lm((log10(Dilutions))~OD$V1)
```

```
> Reciprocal_Regression=lm((1/Dilutions)~OD$V1)
>
> summary(Regression)

Call:
lm(formula = Dilutions ~ OD$V1)

Residuals:
   Min     1Q Median     3Q    Max
-27418  -7277   1505   7155  46693

Coefficients:
            Estimate Std. Error t value Pr(>|t|)
(Intercept)    69634      14464   4.814 0.000541 ***
OD$V1        -130719      32252  -4.053 0.001905 **
---
Signif. codes:  0 ‘***’ 0.001 ‘**’ 0.01 ‘*’ 0.05 ‘.’ 0.1 ‘ ’ 1

Residual standard error: 19200 on 11 degrees of freedom
Multiple R-squared:  0.5989,  Adjusted R-squared:  0.5625
F-statistic: 16.43 on 1 and 11 DF,  p-value: 0.001905

> Titre_1=coef(Regression)[1] + (cutoff)*coef(Regression)[2]
> Titre_1
(Intercept)
   35536.41
>
> summary(log10_Regression)

Call:
lm(formula = (log10(Dilutions)) ~ OD$V1)

Residuals:
    Min      1Q  Median      3Q     Max
-1.3551 -0.1966  0.1863  0.3321  0.7305

Coefficients:
            Estimate Std. Error t value Pr(>|t|)
(Intercept)   5.3732     0.4586  11.716 1.49e-07 ***
OD$V1        -4.8656     1.0226  -4.758 0.000592 ***
---
Signif. codes:  0 ‘***’ 0.001 ‘**’ 0.01 ‘*’ 0.05 ‘.’ 0.1 ‘ ’ 1

Residual standard error: 0.6088 on 11 degrees of freedom
Multiple R-squared:  0.673,   Adjusted R-squared:  0.6433
F-statistic: 22.64 on 1 and 11 DF,  p-value: 0.0005921

> A=coef(log10_Regression)[1] + (cutoff)*coef(log10_Regression)[2]
> Titre_2=10^A
> Titre_2
(Intercept)
   12707.69
>
> summary(Reciprocal_Regression)
```

```
Call:
lm(formula = (1/Dilutions) ~ OD$V1)

Residuals:
       Min         1Q     Median         3Q        Max 
-0.0037382 -0.0032777 -0.0014601 -0.0002329  0.0163973 

Coefficients:
              Estimate Std. Error t value Pr(>|t|)
(Intercept) -0.0009727  0.0043757  -0.222    0.828
OD$V1        0.0095991  0.0097568   0.984    0.346

Residual standard error: 0.005808 on 11 degrees of freedom
Multiple R-squared:  0.08088, Adjusted R-squared:  -0.002679 
F-statistic: 0.9679 on 1 and 11 DF,  p-value: 0.3463

> B=coef(Reciprocal_Regression)[1] + (cutoff)*coef(Reciprocal_Regression)[2]
> Titre_3=1/B
> Titre_3
(Intercept) 
    653.106 

#934-P1
> OD=read.table("clipboard")
> OD
        V1
1  0.57905
2  0.57820
3  0.49925
4  0.54260
5  0.53120
6  0.55375
7  0.52145
8  0.46950
9  0.38430
10 0.38200
11 0.30765
12 0.20630
13 0.14420
> Dilutions=c(50,100,250,500,750,1000,2000,4000,8000,10000,20000,50000,100000)
> 
> #Different approaches to get significant p-value and Rsquare >0.7
> Regression=lm(Dilutions~OD$V1)
> log10_Regression=lm((log10(Dilutions))~OD$V1)
> Reciprocal_Regression=lm((1/Dilutions)~OD$V1)
> 
> summary(Regression)

Call:
lm(formula = Dilutions ~ OD$V1)

Residuals:
   Min     1Q Median     3Q    Max 
-18435  -6499   1672   6429  32433 
```

```
Coefficients:
            Estimate Std. Error t value Pr(>|t|)
(Intercept)    93269      13233   7.048 2.13e-05 ***
OD$V1        -178235      28793  -6.190 6.81e-05 ***
---
Signif. codes:  0 ‘***’ 0.001 ‘**’ 0.01 ‘*’ 0.05 ‘.’ 0.1 ‘ ’ 1

Residual standard error: 14320 on 11 degrees of freedom
Multiple R-squared:  0.777,   Adjusted R-squared:  0.7567
F-statistic: 38.32 on 1 and 11 DF,  p-value: 6.809e-05

> Titre_1=coef(Regression)[1] + (cutoff)*coef(Regression)[2]
> Titre_1
(Intercept)
   46776.97
>
> summary(log10_Regression)

Call:
lm(formula = (log10(Dilutions)) ~ OD$V1)

Residuals:
    Min      1Q  Median      3Q     Max
-0.7260 -0.2680  0.1017  0.2867  0.4995

Coefficients:
            Estimate Std. Error t value Pr(>|t|)
(Intercept)   6.2108     0.3837  16.185  5.1e-09 ***
OD$V1        -6.5381     0.8349  -7.831  8.0e-06 ***
---
Signif. codes:  0 ‘***’ 0.001 ‘**’ 0.01 ‘*’ 0.05 ‘.’ 0.1 ‘ ’ 1

Residual standard error: 0.4152 on 11 degrees of freedom
Multiple R-squared:  0.8479,  Adjusted R-squared:  0.8341
F-statistic: 61.32 on 1 and 11 DF,  p-value: 7.999e-06

> A=coef(log10_Regression)[1] + (cutoff)*coef(log10_Regression)[2]
> Titre_2=10^A
> Titre_2
(Intercept)
   32018.12
>
> summary(Reciprocal_Regression)

Call:
lm(formula = (1/Dilutions) ~ OD$V1)

Residuals:
      Min        1Q    Median        3Q       Max
-0.004326 -0.003399 -0.001806  0.001612  0.014170

Coefficients:
             Estimate Std. Error t value Pr(>|t|)
(Intercept) -0.005699   0.004872  -1.170   0.2668
OD$V1        0.019911   0.010602   1.878   0.0871 .
```

```
---
Signif. codes:  0 ‘***’ 0.001 ‘**’ 0.01 ‘*’ 0.05 ‘.’ 0.1 ‘ ’ 1

Residual standard error: 0.005272 on 11 degrees of freedom
Multiple R-squared:  0.2428,  Adjusted R-squared:  0.174
F-statistic: 3.527 on 1 and 11 DF,  p-value: 0.08711

> B=coef(Reciprocal_Regression)[1] + (cutoff)*coef(Reciprocal_Regression)[2]
> Titre_3=1/B
> Titre_3
(Intercept)
  -1977.013

#934-P2
> OD=read.table("clipboard")
> OD
        V1
1  0.41755
2  0.47225
3  0.46350
4  0.48550
5  0.47860
6  0.48260
7  0.45430
8  0.41660
9  0.32445
10 0.31725
11 0.23105
12 0.05350
13 0.05825
> Dilutions=c(50,100,250,500,750,1000,2000,4000,8000,10000,20000,50000,100000)
>
> #Different approaches to get significant p-value and Rsquare >0.7
> Regression=lm(Dilutions~OD$V1)
> log10_Regression=lm((log10(Dilutions))~OD$V1)
> Reciprocal_Regression=lm((1/Dilutions)~OD$V1)
>
> summary(Regression)

Call:
lm(formula = Dilutions ~ OD$V1)

Residuals:
   Min     1Q Median     3Q    Max
-16149 -11887   2561   5559  35261

Coefficients:
            Estimate Std. Error t value Pr(>|t|)
(Intercept)    74377      10264   7.246 1.65e-05 ***
OD$V1        -165453      26466  -6.252 6.25e-05 ***
---
Signif. codes:  0 ‘***’ 0.001 ‘**’ 0.01 ‘*’ 0.05 ‘.’ 0.1 ‘ ’ 1

Residual standard error: 14210 on 11 degrees of freedom
Multiple R-squared:  0.7804,  Adjusted R-squared:  0.7604
```

```
F-statistic: 39.08 on 1 and 11 DF,  p-value: 6.246e-05

> Titre_1=coef(Regression)[1] + (cutoff)*coef(Regression)[2]
> Titre_1
(Intercept)
   31219.46
>
> summary(log10_Regression)

Call:
lm(formula = (log10(Dilutions)) ~ OD$V1)

Residuals:
    Min      1Q  Median      3Q     Max
-1.3159 -0.3342  0.1987  0.3721  0.5819

Coefficients:
            Estimate Std. Error t value Pr(>|t|)
(Intercept)   5.3297     0.4137  12.882 5.59e-08 ***
OD$V1        -5.5439     1.0668  -5.197 0.000296 ***
---
Signif. codes:  0 ‘***’ 0.001 ‘**’ 0.01 ‘*’ 0.05 ‘.’ 0.1 ‘ ’ 1

Residual standard error: 0.5727 on 11 degrees of freedom
Multiple R-squared:  0.7106,  Adjusted R-squared:  0.6843
F-statistic: 27.01 on 1 and 11 DF,  p-value: 0.000296

> A=coef(log10_Regression)[1] + (cutoff)*coef(log10_Regression)[2]
> Titre_2=10^A
> Titre_2
(Intercept)
   7649.176
>
> summary(Reciprocal_Regression)

Call:
lm(formula = (1/Dilutions) ~ OD$V1)

Residuals:
       Min         1Q     Median         3Q        Max
-0.0036414 -0.0030889 -0.0024577  0.0004452  0.0162832

Coefficients:
             Estimate Std. Error t value Pr(>|t|)
(Intercept) -0.001108   0.004163  -0.266    0.795
OD$V1        0.011556   0.010734   1.077    0.305

Residual standard error: 0.005763 on 11 degrees of freedom
Multiple R-squared:  0.09532, Adjusted R-squared:  0.01308
F-statistic: 1.159 on 1 and 11 DF,  p-value: 0.3047

> B=coef(Reciprocal_Regression)[1] + (cutoff)*coef(Reciprocal_Regression)[2]
> Titre_3=1/B
> Titre_3
(Intercept)
```

```
524.6787
```
