## Supplemental file S7 for "Evaluation of tick salivary and midgut extracellular vesicles as anti-tick vaccines in White-tailed deer (*Odocoileus virginianus*)"

**cutoff-and-titer_ELISA.R**
gonzalezj 2024-07-03

| WTD | Titer |
|---|---|
| 924 – P1 | 100144.1 |
| 929 -P1 | 74191.72 |
| 929 – P2 | 74461.46 |
| 934 – P1 | 77072.79 |

```
##Calculation cutoff value using negative controls (non-vaccinated WTD: 923 and 930)
###NOTE: MG-EVs only for 2 plates of 929 and 930WTD
> ctrl_OD=read.table("clipboard")
> ctrl_OD
        V1
1  0.25845
2  0.24785
3  0.14905
4  0.14635
5  0.10545
6  0.11055
7  0.07630
8  0.06865
9  0.06305
10 0.06785
11 0.05475
12 0.05415
13 0.05500
14 0.24855
15 0.21880
16 0.09945
17 0.08570
18 0.07865
19 0.08325
20 0.06225
21 0.05835
22 0.05140
23 0.05870
24 0.04710
25 0.04925
26 0.04790
27 0.27520
28 0.26675
29 0.15905
30 0.12885
31 0.12745
32 0.12515
33 0.08785
34 0.07800
35 0.06080
36 0.08080
37 0.05290
38 0.05695
39 0.06050
> m=mean(ctrl_OD$V1)
> s=sd(ctrl_OD$V1)
> n=length(ctrl_OD$V1)
> df=n-1
```

```
> tcrit=qt(0.05, df=df, lower.tail = FALSE) #95% confidence level
> f=tcrit*(sqrt(1+(1/n)))
> cutoff_value=m+(f*s)
> cutoff_value
[1] 0.2267414

#ELISA titer determination
#924-P1
> cutoff= 0.2267414 #obtained above
> OD=read.table("clipboard")
> OD
        V1
1  0.63680
2  0.62740
3  0.60310
4  0.61110
5  0.60105
6  0.60465
7  0.60655
8  0.58990
9  0.57840
10 0.55310
11 0.46640
12 0.38310
13 0.28665
> Dilutions=c(50,100,250,500,750,1000,2000,4000,8000,10000,20000,50000,100000)
>
> #Different approaches to get significant p-value and Rsquare >0.7
> Regression=lm(Dilutions~OD$V1)
> log10_Regression=lm((log10(Dilutions))~OD$V1)
> Reciprocal_Regression=lm((1/Dilutions)~OD$V1)
>
> summary(Regression)

Call:
lm(formula = Dilutions ~ OD$V1)

Residuals:
     Min       1Q   Median       3Q      Max
-17086.7   -908.6    288.6   1788.5  15618.7

Coefficients:
            Estimate Std. Error t value Pr(>|t|)
(Intercept)   159803      12399   12.89 5.57e-08 ***
OD$V1        -263113      22171  -11.87 1.30e-07 ***
---
Signif. codes:  0 ‘***’ 0.001 ‘**’ 0.01 ‘*’ 0.05 ‘.’ 0.1 ‘ ’ 1

Residual standard error: 8160 on 11 degrees of freedom
Multiple R-squared:  0.9276,  Adjusted R-squared:  0.921
F-statistic: 140.8 on 1 and 11 DF,  p-value: 1.304e-07

> Titre_1=coef(Regression)[1] + (cutoff)*coef(Regression)[2]
> Titre_1
(Intercept)
```

```
   100144.1
>
> summary(log10_Regression)

Call:
lm(formula = (log10(Dilutions)) ~ OD$V1)

Residuals:
     Min       1Q   Median       3Q      Max
-0.94229 -0.47318  0.00585  0.41512  0.78951

Coefficients:
            Estimate Std. Error t value Pr(>|t|)
(Intercept)   7.7915     0.8699   8.957  2.2e-06 ***
OD$V1        -8.0877     1.5555  -5.199 0.000295 ***
---
Signif. codes:  0 '***' 0.001 '**' 0.01 '*' 0.05 '.' 0.1 ' ' 1

Residual standard error: 0.5725 on 11 degrees of freedom
Multiple R-squared:  0.7108,	Adjusted R-squared:  0.6845
F-statistic: 27.03 on 1 and 11 DF,  p-value: 0.0002947

> A=coef(log10_Regression)[1] + (cutoff)*coef(log10_Regression)[2]
> Titre_2=10^A
> Titre_2
(Intercept)
   907210.9
>
> summary(Reciprocal_Regression)

Call:
lm(formula = (1/Dilutions) ~ OD$V1)

Residuals:
       Min         1Q     Median         3Q        Max
-0.0037872 -0.0032450 -0.0023881  0.0006888  0.0150419

Coefficients:
             Estimate Std. Error t value Pr(>|t|)
(Intercept) -0.009166   0.008412  -1.090    0.299
OD$V1        0.022179   0.015041   1.475    0.168

Residual standard error: 0.005536 on 11 degrees of freedom
Multiple R-squared:  0.165,	Adjusted R-squared:  0.08914
F-statistic: 2.174 on 1 and 11 DF,  p-value: 0.1684

> B=coef(Reciprocal_Regression)[1] + (cutoff)*coef(Reciprocal_Regression)[2]
> Titre_3=1/B
> Titre_3
(Intercept)
  -241.7373

#929-P1
> cutoff= 0.2267414 #obtained above
> OD=read.table("clipboard")
```

```
> OD
        V1
1  0.47205
2  0.50035
3  0.54555
4  0.54445
5  0.54740
6  0.55955
7  0.53380
8  0.51110
9  0.48755
10 0.45165
11 0.36390
12 0.30015
13 0.20880
> Dilutions=c(50,100,250,500,750,1000,2000,4000,8000,10000,20000,50000,100000)
>
> #Different approaches to get significant p-value and Rsquare >0.7
> Regression=lm(Dilutions~OD$V1)
> log10_Regression=lm((log10(Dilutions))~OD$V1)
> Reciprocal_Regression=lm((1/Dilutions)~OD$V1)
>
> summary(Regression)

Call:
lm(formula = Dilutions ~ OD$V1)

Residuals:
     Min       1Q   Median       3Q      Max
-19983.3  -5883.0    729.5   5571.6  21333.6

Coefficients:
            Estimate Std. Error t value Pr(>|t|)
(Intercept)   130743      14195   9.210 1.67e-06 ***
OD$V1        -249408      29881  -8.347 4.35e-06 ***
---
Signif. codes:  0 ‘***’ 0.001 ‘**’ 0.01 ‘*’ 0.05 ‘.’ 0.1 ‘ ’ 1

Residual standard error: 11200 on 11 degrees of freedom
Multiple R-squared:  0.8636, Adjusted R-squared:  0.8512
F-statistic: 69.67 on 1 and 11 DF,  p-value: 4.354e-06

> Titre_1=coef(Regression)[1] + (cutoff)*coef(Regression)[2]
> Titre_1
(Intercept)
   74191.72
>
> summary(log10_Regression)

Call:
lm(formula = (log10(Dilutions)) ~ OD$V1)

Residuals:
    Min      1Q  Median      3Q     Max
-1.5856 -0.1398  0.2030  0.4516  0.7278
```

```
Coefficients:
            Estimate Std. Error t value Pr(>|t|)
(Intercept)   6.6112     0.8962   7.377  1.4e-05 ***
OD$V1        -7.0473     1.8864  -3.736  0.00329 **
---
Signif. codes:  0 ‘***’ 0.001 ‘**’ 0.01 ‘*’ 0.05 ‘.’ 0.1 ‘ ’ 1

Residual standard error: 0.7068 on 11 degrees of freedom
Multiple R-squared:  0.5592,  Adjusted R-squared:  0.5192
F-statistic: 13.96 on 1 and 11 DF,  p-value: 0.00329

> A=coef(log10_Regression)[1] + (cutoff)*coef(log10_Regression)[2]
> Titre_2=10^A
> Titre_2
(Intercept)
   103117.4
>
> summary(Reciprocal_Regression)

Call:
lm(formula = (1/Dilutions) ~ OD$V1)

Residuals:
       Min         1Q     Median         3Q        Max
-0.0032110 -0.0029004 -0.0020760 -0.0007094  0.0168931

Coefficients:
             Estimate Std. Error t value Pr(>|t|)
(Intercept) -0.001174   0.007571  -0.155    0.880
OD$V1        0.009069   0.015937   0.569    0.581

Residual standard error: 0.005971 on 11 degrees of freedom
Multiple R-squared:  0.0286,  Adjusted R-squared:  -0.05971
F-statistic: 0.3238 on 1 and 11 DF,  p-value: 0.5808

> B=coef(Reciprocal_Regression)[1] + (cutoff)*coef(Reciprocal_Regression)[2]
> Titre_3=1/B
> Titre_3
(Intercept)
   1133.673

#929-P2
> cutoff= 0.2267414 #obtained above
> OD=read.table("clipboard")
> OD
        V1
1  0.55160
2  0.57515
3  0.66485
4  0.71150
5  0.70440
6  0.71185
7  0.68700
8  0.65790
```

```
9  0.56645
10 0.58525
11 0.49070
12 0.29925
13 0.23965
> Dilutions=c(50,100,250,500,750,1000,2000,4000,8000,10000,20000,50000,100000)
>
> #Different approaches to get significant p-value and Rsquare >0.7
> Regression=lm(Dilutions~OD$V1)
> log10_Regression=lm((log10(Dilutions))~OD$V1)
> Reciprocal_Regression=lm((1/Dilutions)~OD$V1)
>
> summary(Regression)

Call:
lm(formula = Dilutions ~ OD$V1)

Residuals:
   Min     1Q Median     3Q    Max
-18701  -9195    920   8202  27752

Coefficients:
            Estimate Std. Error t value Pr(>|t|)
(Intercept)   113345      14891   7.612 1.04e-05 ***
OD$V1        -171491      25192  -6.807 2.92e-05 ***
---
Signif. codes:  0 ‘***’ 0.001 ‘**’ 0.01 ‘*’ 0.05 ‘.’ 0.1 ‘ ’ 1

Residual standard error: 13280 on 11 degrees of freedom
Multiple R-squared:  0.8082,  Adjusted R-squared:  0.7907
F-statistic: 46.34 on 1 and 11 DF,  p-value: 2.925e-05

> Titre_1=coef(Regression)[1] + (cutoff)*coef(Regression)[2]
> Titre_1
(Intercept)
   74461.46
>
> summary(log10_Regression)

Call:
lm(formula = (log10(Dilutions)) ~ OD$V1)

Residuals:
     Min       1Q   Median       3Q      Max
-1.73926 -0.02933  0.17678  0.53079  0.71117

Coefficients:
            Estimate Std. Error t value Pr(>|t|)
(Intercept)   5.8872     0.8939   6.586 3.94e-05 ***
OD$V1        -4.4398     1.5123  -2.936   0.0135 *
---
Signif. codes:  0 ‘***’ 0.001 ‘**’ 0.01 ‘*’ 0.05 ‘.’ 0.1 ‘ ’ 1

Residual standard error: 0.7972 on 11 degrees of freedom
Multiple R-squared:  0.4393,  Adjusted R-squared:  0.3883
```

```
F-statistic: 8.619 on 1 and 11 DF,  p-value: 0.01355

> A=coef(log10_Regression)[1] + (cutoff)*coef(log10_Regression)[2]
> Titre_2=10^A
> Titre_2
(Intercept)
   75950.99
>
> summary(Reciprocal_Regression)

Call:
lm(formula = (1/Dilutions) ~ OD$V1)

Residuals:
      Min        1Q    Median        3Q       Max
-0.002972 -0.002794 -0.002344 -0.001344  0.017018

Coefficients:
            Estimate Std. Error t value Pr(>|t|)
(Intercept) 0.001735   0.006782   0.256    0.803
OD$V1       0.002261   0.011474   0.197    0.847

Residual standard error: 0.006048 on 11 degrees of freedom
Multiple R-squared:  0.003516,	Adjusted R-squared:  -0.08707
F-statistic: 0.03882 on 1 and 11 DF,  p-value: 0.8474

> B=coef(Reciprocal_Regression)[1] + (cutoff)*coef(Reciprocal_Regression)[2]
> Titre_3=1/B
> Titre_3
(Intercept)
   444.8921

#934-P1
> cutoff= 0.2267414 #obtained above
> OD=read.table("clipboard")
> OD
        V1
1  0.52680
2  0.53090
3  0.54705
4  0.51620
5  0.54325
6  0.55815
7  0.56535
8  0.52630
9  0.43240
10 0.45630
11 0.40600
12 0.30005
13 0.21830
> Dilutions=c(50,100,250,500,750,1000,2000,4000,8000,10000,20000,50000,100000)
>
> #Different approaches to get significant p-value and Rsquare >0.7
> Regression=lm(Dilutions~OD$V1)
> log10_Regression=lm((log10(Dilutions))~OD$V1)
```

```
> Reciprocal_Regression=lm((1/Dilutions)~OD$V1)
>
> summary(Regression)

Call:
lm(formula = Dilutions ~ OD$V1)

Residuals:
     Min       1Q   Median       3Q      Max
-16982.6  -8504.8     65.9   4306.5  20789.1

Coefficients:
            Estimate Std. Error t value Pr(>|t|)
(Intercept)   134503      13762   9.774 9.29e-07 ***
OD$V1        -253285      28524  -8.880 2.39e-06 ***
---
Signif. codes:  0 '***' 0.001 '**' 0.01 '*' 0.05 '.' 0.1 ' ' 1

Residual standard error: 10610 on 11 degrees of freedom
Multiple R-squared:  0.8776,  Adjusted R-squared:  0.8664
F-statistic: 78.85 on 1 and 11 DF,  p-value: 2.391e-06

> Titre_1=coef(Regression)[1] + (cutoff)*coef(Regression)[2]
> Titre_1
(Intercept)
   77072.79
>
> summary(log10_Regression)

Call:
lm(formula = (log10(Dilutions)) ~ OD$V1)

Residuals:
     Min       1Q   Median       3Q      Max
-1.21593 -0.30275  0.08749  0.45111  0.68451

Coefficients:
            Estimate Std. Error t value Pr(>|t|)
(Intercept)   6.9924     0.7998   8.743 2.78e-06 ***
OD$V1        -7.7402     1.6577  -4.669 0.000683 ***
---
Signif. codes:  0 '***' 0.001 '**' 0.01 '*' 0.05 '.' 0.1 ' ' 1

Residual standard error: 0.6165 on 11 degrees of freedom
Multiple R-squared:  0.6647,  Adjusted R-squared:  0.6342
F-statistic:  21.8 on 1 and 11 DF,  p-value: 0.0006833

> A=coef(log10_Regression)[1] + (cutoff)*coef(log10_Regression)[2]
> Titre_2=10^A
> Titre_2
(Intercept)
   172748.9
>
> summary(Reciprocal_Regression)
```

```
Call:
lm(formula = (1/Dilutions) ~ OD$V1)

Residuals:
       Min         1Q     Median         3Q        Max
-0.0041115 -0.0029065 -0.0018814 -0.0001294  0.0160369

Coefficients:
             Estimate Std. Error t value Pr(>|t|)
(Intercept) -0.004897   0.007469  -0.656    0.526
OD$V1        0.016819   0.015481   1.086    0.301

Residual standard error: 0.005758 on 11 degrees of freedom
Multiple R-squared:  0.0969,  Adjusted R-squared:  0.0148
F-statistic:  1.18 on 1 and 11 DF,  p-value: 0.3005

> B=coef(Reciprocal_Regression)[1] + (cutoff)*coef(Reciprocal_Regression)[2]
> Titre_3=1/B
> Titre_3
(Intercept)
  -922.8738
```
