## Supplemental file S12 for "Evaluation of tick salivary and midgut extracellular vesicles as anti-tick vaccines in White-tailed deer (*Odocoileus virginianus*)"

**EV_nymphs.R**

gonzalezj 2024-07-10

```r
##NYMPH TICKS INFESTATION ON DEER_VACCINE PROJECT TAMU##
View(nymphs_deer)
#comparing EN recovered ALIVE
table_survival <- table(nymphs_deer$treatment, nymphs_deer$EN_survival)
chisq.test(table_survival)

##
##  Pearson's Chi-squared test with Yates' continuity correction
##
## data:  table_survival
## X-squared = 15.355, df = 1, p-value = 8.911e-05

print(table_survival)

##
##                 0   1
##   control      25 151
##   vaccinated    3 159

#comparing EN molted from EN survived
table_molted <- table(nymphs_deer$treatment, nymphs_deer$EN_molted)
chisq.test(table_molted)

##
##  Pearson's Chi-squared test with Yates' continuity correction
##
## data:  table_molted
## X-squared = 13.955, df = 1, p-value = 0.0001873

print(table_molted)

##
##                 0   1
##   control      34 117
##   vaccinated   11 148

weights <- na.omit(nymphs_deer)

# Perform Shapiro-Wilk test for EN_weight
weights %>%
  group_by(treatment) %>%
  summarise(shapiro_test_p_value = shapiro.test(EN_weight)$p.value)

## # A tibble: 2 × 2
##   treatment  shapiro_test_p_value
##   <fct>                     <dbl>
## 1 control                   0.777
## 2 vaccinated                0.00580

transformed <- weights %>%
  mutate(log_EN= log(EN_weight + 0.2),
         sqrt_EN = sqrt(EN_weight))
```

```r
# Performing Shapiro-Wilk test after log transformation
transformed %>%
  group_by(treatment) %>%
  summarise(
    n = n(),
    shapiro_test_p_value_log = ifelse(n >= 3 && n <= 5000, shapiro.test(log_EN)$p.value,
NA)
  ) %>%
  filter(!is.na(shapiro_test_p_value_log))

## # A tibble: 2 × 3
##   treatment      n shapiro_test_p_value_log
##   <fct>      <int>                    <dbl>
## 1 control       60                  0.825
## 2 vaccinated    60                  0.00529

###LOG TRANSFORMATION NOT WORKING###

# Performing Shapiro-Wilk test after square root transformation
transformed %>%
  group_by(treatment) %>%
  summarise(
    n = n(),
    shapiro_test_p_value_sqrt = ifelse(n >= 3 && n <= 5000,
shapiro.test(sqrt_EN)$p.value, NA)
  ) %>%
  filter(!is.na(shapiro_test_p_value_sqrt))

## # A tibble: 2 × 3
##   treatment      n shapiro_test_p_value_sqrt
##   <fct>      <int>                     <dbl>
## 1 control       60                   0.00776
## 2 vaccinated    60                   0.00174

###SQRT TRANSFORMATION NOT WORKING###

#Performing non-parametric Wilcoxon test for 2 groups
wilcox_EN <- wilcox.test(EN_weight ~ treatment, data = weights)
print(wilcox_EN)

##
##  Wilcoxon rank sum test with continuity correction
##
## data:  EN_weight by treatment
## W = 1694.5, p-value = 0.5815
## alternative hypothesis: true location shift is not equal to 0

W_summary <- weights %>%
  group_by(treatment) %>%
  summarise(
    mean_value = mean(EN_weight),
    sem_value = sd(EN_weight) / sqrt(n()),
    ci_low = mean(EN_weight) - qt(0.975, n()-1) * sem_value,
    ci_high = mean(EN_weight) + qt(0.975, n()-1) * sem_value,
    .groups='drop'
```

```
  )
print(W_summary)

## # A tibble: 2 × 5
##   treatment  mean_value sem_value  ci_low ci_high
##   <fct>           <dbl>     <dbl>   <dbl>   <dbl>
## 1 control       0.00766  0.000415 0.00683 0.00849
## 2 vaccinated    0.00798  0.000341 0.00729 0.00866
```

EV_females.R

```
##FEMALE TICK INFESTATION_EV-VACCINE ON WTD##
#Fisher test to compare the results
View(EF_deer)
table_attached <- table(EF_deer$treatment, EF_deer$attached)
print(table_attached)

## 
##              EF_dead_feeding EF_dead_squished
##   control                  0                5
##   vaccinated              21                8

test_attached <- fisher.test(table_attached, conf.int = TRUE)
print(test_attached)

## 
##  Fisher's Exact Test for Count Data
## 
## data:  table_attached
## p-value = 0.004625
## alternative hypothesis: true odds ratio is not equal to 1
## 95 percent confidence interval:
##  0.0000000 0.5430005
## sample estimates:
## odds ratio 
##          0

table_detached <- table(EF_deer$treatment, EF_deer$detached)
print(table_detached)

## 
##              EF_dead_post EF_oviposition
##   control              16             18
##   vaccinated           16             32

test_detached <- fisher.test(table_detached, conf.int = TRUE)
print(test_detached)

## 
##  Fisher's Exact Test for Count Data
## 
## data:  table_detached
## p-value = 0.2538
## alternative hypothesis: true odds ratio is not equal to 1
## 95 percent confidence interval:
```

```
##  0.6550656 4.8179215
## sample estimates:
## odds ratio
##   1.765103

# Collecting p-values from tests
p_values <- c(test_attached$p.value, test_detached$p.value)

# Adjust p-values using Holm-Bonferroni method
p_adjusted <- p.adjust(p_values, method = "holm")
print(p_adjusted)

## [1] 0.009250474 0.253786825

View(EFovi_deer)
# Perform Shapiro-Wilk test NORMALITY for EF_weight
EFovi_deer %>%
  group_by(treatment) %>%
  summarise(shapiro_test_p_value = shapiro.test(EF_weight)$p.value)

## # A tibble: 2 × 2
##   treatment  shapiro_test_p_value
##   <fct>                     <dbl>
## 1 control                  0.0210
## 2 vaccinated               0.00550

transformedEF <- EFovi_deer %>%
  mutate(log_EF= log(EF_weight + 0.2),
         sqrt_EF = sqrt(EF_weight))
transformedEF %>%
  group_by(treatment) %>%
  summarise(
    n = n(),
    shapiro_test_p_value_log = ifelse(n >= 3 && n <= 5000, shapiro.test(log_EF)$p.value,
NA)
  ) %>%
  filter(!is.na(shapiro_test_p_value_log))

## # A tibble: 2 × 3
##   treatment      n shapiro_test_p_value_log
##   <fct>      <int>                    <dbl>
## 1 control       33                0.00179
## 2 vaccinated    48                0.0000625

###LOG TRANSFORMATION NOT WORKING###

# Performing Shapiro-Wilk test after square root transformation
transformedEF %>%
  group_by(treatment) %>%
  summarise(
    n = n(),
    shapiro_test_p_value_sqrt = ifelse(n >= 3 && n <= 5000,
shapiro.test(sqrt_EF)$p.value, NA)
  ) %>%
  filter(!is.na(shapiro_test_p_value_sqrt))
```

```
## # A tibble: 2 × 3
##   treatment      n shapiro_test_p_value_sqrt
##   <fct>      <int>                     <dbl>
## 1 control       33                  0.00418
## 2 vaccinated    48                  0.000143

###SQRT TRANSFORMATION NOT WORKING###

#Performing non-parametric Wilcoxon test for 2 groups
wilcox_EF <- wilcox.test(EF_weight ~ treatment, data = EFovi_deer)
print(wilcox_EF)

##
##  Wilcoxon rank sum exact test
##
## data:  EF_weight by treatment
## W = 885, p-value = 0.376
## alternative hypothesis: true location shift is not equal to 0

EF_summary <- EFovi_deer %>%
  group_by(treatment) %>%
  summarise(
    mean_value = mean(EF_weight),
    sem_value = sd(EF_weight) / sqrt(n()),
    ci_low = mean(EF_weight) - qt(0.975, n()-1) * sem_value,
    ci_high = mean(EF_weight) + qt(0.975, n()-1) * sem_value,
    .groups='drop'
  )
print(EF_summary)

## # A tibble: 2 × 5
##   treatment  mean_value sem_value ci_low ci_high
##   <fct>           <dbl>     <dbl>  <dbl>   <dbl>
## 1 control         0.402    0.0472  0.306   0.498
## 2 vaccinated      0.360    0.0312  0.297   0.423

# Perform Shapiro-Wilk test NORMALITY for eggs_weight
Eweight <- EFovi_deer[!is.na(EFovi_deer$eggs_weight), ]
Eweight %>%
  group_by(treatment) %>%
  summarise(shapiro_test_p_value = shapiro.test(eggs_weight)$p.value)

## # A tibble: 2 × 2
##   treatment  shapiro_test_p_value
##   <fct>                     <dbl>
## 1 control                   0.454
## 2 vaccinated                0.741

#Differences between groups normally distributed: Welch's correction
t.test(eggs_weight ~ treatment, data = Eweight, var.equal = FALSE)

##
##  Welch Two Sample t-test
##
## data:  eggs_weight by treatment
## t = 0.11044, df = 25.254, p-value = 0.9129
## alternative hypothesis: true difference in means between group control and group
```

```
vaccinated is not equal to 0
## 95 percent confidence interval:
##  -0.06802650  0.07573969
## sample estimates:
##    mean in group control mean in group vaccinated
##                0.2341722                0.2303156

Egg_summary <- Eweight %>%
  group_by(treatment) %>%
  summarise(
    mean_value = mean(eggs_weight),
    sem_value = sd(eggs_weight) / sqrt(n()),
    ci_low = mean(eggs_weight) - qt(0.975, n()-1) * sem_value,
    ci_high = mean(eggs_weight) + qt(0.975, n()-1) * sem_value,
    .groups='drop'
  )
print(Egg_summary)

## # A tibble: 2 × 5
##   treatment  mean_value sem_value ci_low ci_high
##   <fct>           <dbl>     <dbl>  <dbl>   <dbl>
## 1 control         0.234    0.0314  0.168   0.300
## 2 vaccinated      0.230    0.0153  0.199   0.262

# Perform Shapiro-Wilk test NORMALITY for estimate LARVA MASS
larvae <- EFovi_deer[!is.na(EFovi_deer$hatch_estL), ]
larvae %>%
  group_by(treatment) %>%
  summarise(shapiro_test_p_value = shapiro.test(hatch_estL)$p.value)

## # A tibble: 2 × 2
##   treatment  shapiro_test_p_value
##   <fct>                     <dbl>
## 1 control                   0.149
## 2 vaccinated                0.722

#Differences between groups normally distributed: Welch's correction
t.test(hatch_estL ~ treatment, data = larvae, var.equal = FALSE)

##
##  Welch Two Sample t-test
##
## data:  hatch_estL by treatment
## t = -0.11095, df = 26.676, p-value = 0.9125
## alternative hypothesis: true difference in means between group control and group
vaccinated is not equal to 0
## 95 percent confidence interval:
##  -1057.259   948.845
## sample estimates:
##    mean in group control mean in group vaccinated
##                 2444.000                 2498.207

L_summary <- larvae %>%
  group_by(treatment) %>%
  summarise(
    mean_value = mean(hatch_estL),
    sem_value = sd(hatch_estL) / sqrt(n()),
```

```
    ci_low = mean(hatch_estL) - qt(0.975, n()-1) * sem_value,
    ci_high = mean(hatch_estL) + qt(0.975, n()-1) * sem_value,
    .groups='drop'
  )
print(L_summary)

## # A tibble: 2 × 5
##   treatment  mean_value sem_value ci_low ci_high
##   <fct>           <dbl>     <dbl>  <dbl>   <dbl>
## 1 control         2444       423.  1546.   3342.
## 2 vaccinated      2498.      244.  1999.   2997.

# Perform Shapiro-Wilk test NORMALITY for %Hatchability
larvae %>%
  group_by(treatment) %>%
  summarise(shapiro_test_p_value = shapiro.test(hatch_percent)$p.value)

## # A tibble: 2 × 2
##   treatment  shapiro_test_p_value
##   <fct>                     <dbl>
## 1 control               0.000608
## 2 vaccinated            0.0000355

transformedHatch <- larvae %>%
  mutate(log_H= log(hatch_percent + 0.2),
         sqrt_H = sqrt(hatch_percent))
transformedHatch %>%
  group_by(treatment) %>%
  summarise(
    n = n(),
    shapiro_test_p_value_log = ifelse(n >= 3 && n <= 5000, shapiro.test(log_H)$p.value,
NA)
  ) %>%
  filter(!is.na(shapiro_test_p_value_log))

## # A tibble: 2 × 3
##   treatment      n shapiro_test_p_value_log
##   <fct>      <int>                    <dbl>
## 1 control       17             0.000000602
## 2 vaccinated    29             0.00000000184

###LOG TRANSFORMATION NOT WORKING###

# Performing Shapiro-Wilk test after square root transformation
transformedHatch  %>%
  group_by(treatment) %>%
  summarise(
    n = n(),
    shapiro_test_p_value_sqrt = ifelse(n >= 3 && n <= 5000, shapiro.test(sqrt_H)$p.value,
NA)
  ) %>%
  filter(!is.na(shapiro_test_p_value_sqrt))

## # A tibble: 2 × 3
##   treatment      n shapiro_test_p_value_sqrt
##   <fct>      <int>                     <dbl>
```

```
## 1 control       17                0.0000275
## 2 vaccinated    29                0.000000229

###SQRT TRANSFORMATION NOT WORKING###

#Performing non-parametric Wilcoxon test for 2 groups
wilcox_H <- wilcox.test(hatch_percent ~ treatment, data = larvae)

## Warning in wilcox.test.default(x = DATA[[1L]], y = DATA[[2L]], ...): cannot
## compute exact p-value with ties

print(wilcox_H)

##
##  Wilcoxon rank sum test with continuity correction
##
## data:  hatch_percent by treatment
## W = 274.5, p-value = 0.5314
## alternative hypothesis: true location shift is not equal to 0

Hatch_summary <- larvae %>%
  group_by(treatment) %>%
  summarise(
    mean_value = mean(hatch_percent),
    sem_value = sd(hatch_percent) / sqrt(n()),
    ci_low = mean(hatch_percent) - qt(0.975, n()-1) * sem_value,
    ci_high = mean(hatch_percent) + qt(0.975, n()-1) * sem_value,
    .groups='drop'
  )
print(Hatch_summary)

## # A tibble: 2 × 5
##   treatment  mean_value sem_value ci_low ci_high
##   <fct>           <dbl>     <dbl>  <dbl>   <dbl>
## 1 control          77.9      6.58   63.9    91.8
## 2 vaccinated       74.9      5.04   64.6    85.2
```

```
> View(EFovi_deer)
> oviposition_time <- EFovi_deer[!is.na(EFovi_deer$oviposition_time), ]
> oviposition_time %>%
+   group_by(treatment) %>%
+   summarise(shapiro_test_p_value = shapiro.test(oviposition_time)$p.value)
# A tibble: 2 × 2
  treatment  shapiro_test_p_value
  <fct>                     <dbl>
1 control                  0.0543
2 vaccinated               0.0396
> View(oviposition_time)
> transformedovi <- oviposition_time %>%
+   mutate(log_ovi= log(oviposition_time + 0.2),
+          sqrt_ovi = sqrt(oviposition_time))
> transformedovi %>%
+   group_by(treatment) %>%
+   summarise(
+     n = n(),
+     shapiro_test_p_value_log = ifelse(n >= 3 && n <= 5000, shapiro.test(log_ovi)$p.value, NA)
+   ) %>%
+   filter(!is.na(shapiro_test_p_value_log))
# A tibble: 2 × 3
  treatment      n shapiro_test_p_value_log
  <fct>      <int>                    <dbl>
1 control       18                    0.405
2 vaccinated    32                    0.130
> View(transformedovi)
> #Differences between groups normally distributed: Welch's correction
> t.test(log_ovi ~ treatment, data = transformedovi, var.equal = FALSE)

        Welch Two Sample t-test

data:  log_ovi by treatment
t = -1.8407, df = 33.678, p-value = 0.0745
alternative hypothesis: true difference in means between group control and group vaccinated is not equal to 0
95 percent confidence interval:
 -0.21447712  0.01064779
sample estimates:
   mean in group control mean in group vaccinated
                3.602880                 3.704794

> Ovi_summary <- oviposition_time %>%
+   group_by(treatment) %>%
+   summarise(
+     mean_value = mean(oviposition_time),
+     sem_value = sd(oviposition_time) / sqrt(n()),
+     ci_low = mean(oviposition_time) - qt(0.975, n()-1) * sem_value,
+     ci_high = mean(oviposition_time) + qt(0.975, n()-1) * sem_value,
+     .groups='drop'
+   )
> print(Ovi_summary)
# A tibble: 2 × 5
  treatment  mean_value sem_value ci_low ci_high
  <fct>           <dbl>     <dbl>  <dbl>   <dbl>
1 control          37.2      1.78   33.4    40.9
2 vaccinated       41.1      1.33   38.4    43.8
```
